# Genome-wide CRISPR and small-molecule screens uncover targetable dependencies in ATRT

**DOI:** 10.1101/2020.12.09.417378

**Authors:** Daniel J. Merk, Sophie Hirsch, Foteini Tsiami, Bianca Walter, Lara A. Haeusser, Sepideh Babaei, Jakob Admar, Nicolas Casadei, Cristiana Roggia, Michael Spohn, Jens Schittenhelm, Stephan Singer, Ulrich Schüller, Federica Piccioni, Nicole S. Persky, David E. Root, Manfred Claassen, Marcos Tatagiba, Ghazaleh Tabatabai

**Affiliations:** Department of Neurology and Interdisciplinary Neuro-Oncology; Hertie Institute for Clinical Brain Research; University Hospital Tübingen; Eberhard Karls University Tübingen; Tübingen, 72076; Germany; Internal Medicine I; University Hospital Tübingen; Eberhard Karls University Tübingen; Tübingen, 72076; Germany; Institute of Medical Genetics and Applied Genomics; University Hospital Tübingen; Eberhard Karls University Tübingen; Tübingen, 72076; Germany; NGS Competence Center Tübingen; Eberhard Karls University Tübingen; Tübingen, 72076; Germany; Research Institute Children’s Cancer Center; Hamburg, 20251; Germany; Bioinformatics Core; University Medical Center Hamburg-Eppendorf; Hamburg, 20251; Germany; Institute of Pathology and Neuropathology, Department of Neuropathology; University Hospital Tübingen; Eberhard Karls University Tübingen; Tübingen, 72076; Germany; Institute of Pathology and Neuropathology; Department of General and Molecular Pathology, University Hospital Tübingen; Eberhard Karls University Tübingen; Tübingen, 72076; Germany; Department of Pediatric Hematology and Oncology; University Hospital Hamburg-Eppendorf; Hamburg, 20246; Germany; Institute of Neuropathology; University Hospital Hamburg-Eppendorf; Hamburg, 20246; Germany; Genetic Perturbation Platform; Broad Institute of MIT and Harvard; Cambridge, MA, 02142; USA; Merck Research Laboratories; Boston, MA, 02115; USA; Department of Computer Science, Eberhard Karls University Tübingen; Tübingen, 72076, Germany; Department of Neurosurgery; University Hospital Tübingen; Eberhard Karls University Tübingen; Tübingen, 72076, Germany; Cluster of Excellence iFIT (EXC 2180) “Image Guided and Functionally Instructed Tumor Therapies”; Eberhard Karls University Tübingen; Tübingen, 72076; Germany; German Consortium for Translational Cancer Research (DKTK); Partner Site Tübingen; German Cancer Research Center (DKFZ); Heidelberg, 69120; Germany; Cluster of Excellence (EXC 2064) “Machine learning”, Eberhard Karls University Tübingen; Tübingen, 72076; Germany; Comprehensive Cancer Center Tübingen Stuttgart, University Hospital Tübingen, Eberhard Karls University Tübingen, Tübingen, 72076; Germany

## Abstract

Atypical teratoid rhabdoid tumors (ATRT) are incurable high-grade pediatric brain tumors. Concepts for molecular-driven therapies in ATRTs lag behind, mainly due to the absence of actionable genetic alterations. We performed genome-wide CRISPR/Cas9 knockout screens in six human ATRT cell lines and identified a total of 671 context-specific essential genes. Based on these genetic dependencies, we constructed a library of small-molecule inhibitors that we found to preferentially inhibit growth of ATRT cells. CDK4/6 inhibitors, among the most potent drugs in our library, are capable of inhibiting tumor growth due to mutual exclusive dependency of ATRTs on *CDK4* or *CDK6*. These distinct dependencies drive heterogeneity in response to CDK4/6 inhibitors in ATRTs. Our approach might serve as a blueprint for fostering the identification of functionally-instructed therapeutic strategies in other incurable diseases beyond ATRT, whose genomic profiles also lack actionable alterations so far.

## Introduction

Genome-scale perturbation screens are a powerful tool to dissect context-dependent functional networks in cancer cells on a single gene level. Loss-of-function strategies using shRNA or CRISPR/Cas9 techniques have thus been used to reveal genetic dependencies, which can guide the development of functionally-instructed therapies for distinct tumor entities (Ghandi et al., 2019). This approach seems to be especially useful for those diseases, where conventional high-throughput next-generation sequencing strategies failed to identify actionable and druggable molecular alterations. This challenge becomes particularly evident in several types of pediatric brain tumors (Gröbner et al., 2018), the leading cause of cancer-related deaths in children and adolescents.

Atypical teratoid rhabdoid tumors (ATRT) is a highly malignant brain tumor that accounts for up to 50 % of all central nervous system neoplasms in the first year of life (Hilden et al., 2004; Ostrom et al., 2019; Woehrer et al., 2010). Loss-of-function alterations of *SMARCB1*, and rarely *SMARCA4*, serve as diagnostic molecular markers. The presence of alterations in these two components of the SWI/SNF chromatin remodelling complex suggests a major role for epigenetic dysregulation during the initiation and/or progression of ATRTs (Biegel et al., 2002; Hasselblatt et al., 2014; Schneppenheim et al., 2010; Versteege et al., 1998). From a clinical perspective, however, the loss of a tumor suppressor (e.g. *SMARCB1*) gene cannot be exploited as therapeutic target. Thus, novel approaches of molecular profiling are needed.

In spite of their consistent homogeneity on the genetic level, ATRTs display a remarkable epigenetic and transcriptional heterogeneity (Johann et al., 2016; Torchia et al., 2016). Based on these studies, ATRTs are currently segregated into three molecular subgroups: ATRT-SHH (formerly group 1), ATRT-TYR (group 2A), and ATRT-MYC (group 2B) (Ho et al., 2020). Whereas these distinct molecular profiles clearly correlate with clinical features, such as patient age and tumor location, it remains elusive if these molecular subgroups also enable molecular-driven therapeutic strategies.

We hypothesized that genome-wide essentiality screens will identify genetic dependencies and functionally relevant molecular targets in ATRTs despite their “untargetable” genetic profile. Indeed, negative selection CRISPR/Cas9 knockout screens and informed small-molecule drug assays converged to reveal targetable vulnerabilities in ATRTs. These vulnerabilities were independent from molecular subgroups of ATRTs, suggesting a dominant role of the shared *SMARCB1* alteration as a predisposing genetic factor to the discovered set of vulnerabilities. Our data provide a first comprehensive overview of genetic essentialities in ATRTs and reveal clinically available agents such as CDK4/6 inhibitors that might enable rapid translation to improve the outcome of ATRT patients. Most importantly, our study demonstrates the utility of large-scale functional genetic screening and subsequent validation in relevant preclinical models as a powerful platform approach for functionally-instructed target discovery in cancer entities lacking actionable genetic mutations.

## Results

### CRISPR/Cas9 knockout screens reveal genetic dependencies of ATRTs

In preparation for a large-scale perturbational screening approach, we first performed a detailed characterization of a set of seven human ATRT cell lines (BT12, BT16, CHLA02, CHLA04, CHLA05, CHLA06, CHLA266) including genetic, epigenetic, and transcriptional profiling. ATRT cell lines presented rather stable genomes with the exception of the *SMARCB1* locus which was altered in all cell lines by either copy number loss and/or single nucleotide variation (Figure S1A,B; Table S1). No fusion genes were detected by RNA-seq analyses (data not shown). Based on global DNA methylation, ATRT cell lines matched best to primary ATRTs in a reference cohort of > 2,800 CNS tumors (Figure 1A). To further affiliate these cell lines with previously described molecular subgroups of ATRTs (Ho et al., 2020), we employed both gene set enrichment analyses and machine-learning prediction approaches on global transcriptional and DNA methylation profiles, respectively. Three cell lines (CHLA02, CHLA04, CHLA05) were consistently assigned to the ATRT-SHH (group 1) subgroup (Figure S1C-F). The remaining four cell lines (BT12, BT16, CHLA06, CHLA266) displayed a considerable degree of heterogeneity with respect to subgroup-specific epigenetic and transcriptional profiles. Yet, when taking our whole molecular data set into account, we suggest that BT12, BT16, CHLA06, CHLA266 cells fit best to the ATRT-TYR/MYC molecular subgroup (group 2) (Fig. S1G) as has been recently suggested (Ho et al., 2020). Thus, our collection of ATRT cell lines recapitulates the epigenetic and transcriptional diversity of primary ATRTs.

**Figure 1.**
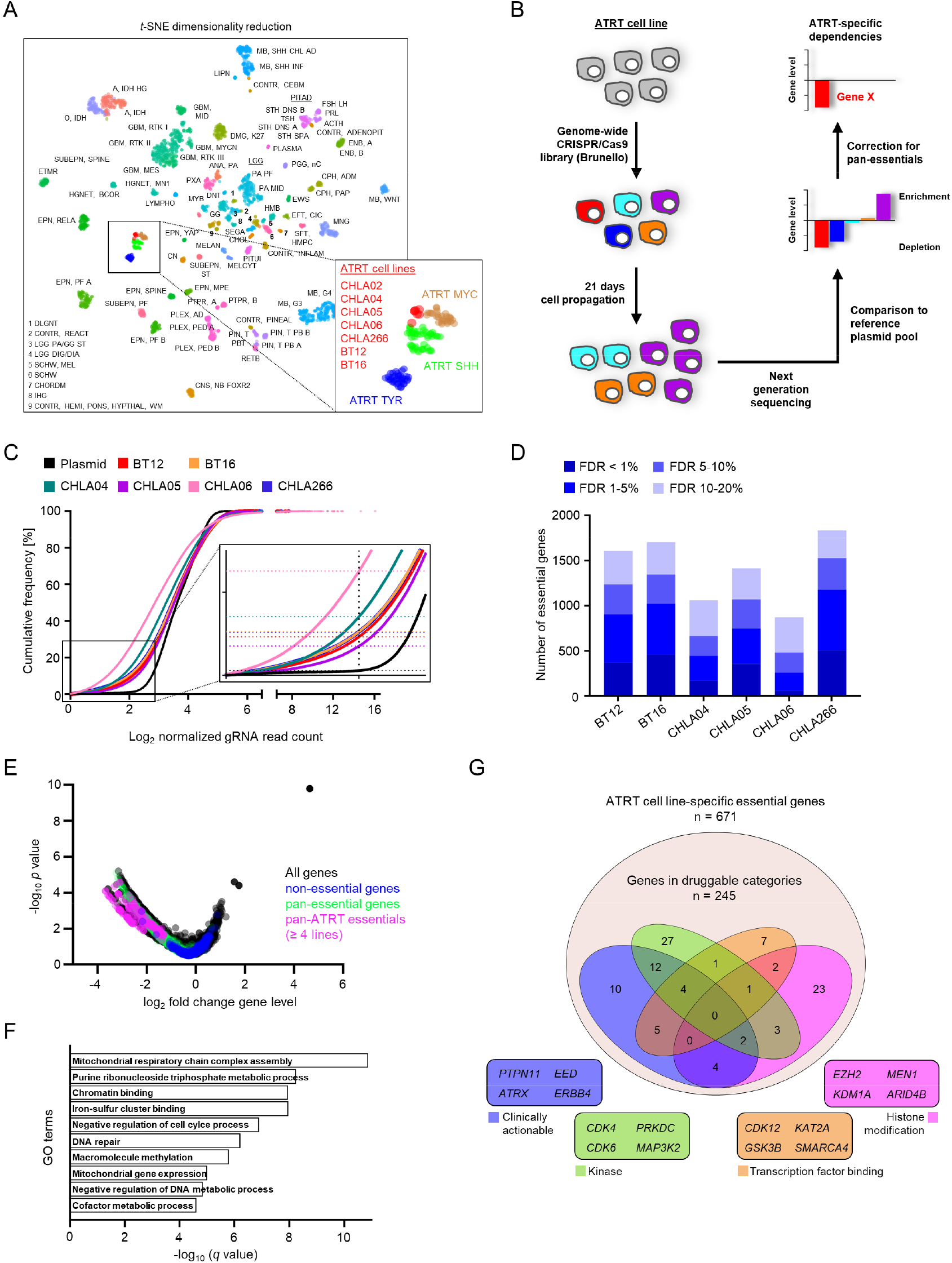
CRISPR/Cas9 knockout screens reveal genetic dependencies of ATRTs. (A) *t*-SNE dimensionality reduction of global DNA methylation profiles from seven human ATRT cell lines (indicated in red) and a reference cohort of 2,801 primary CNS tumors. (B) Overview of experimental approach using CRISPR/Cas9 knockout screens to identify ATRT-specific genetic dependencies. (C) Cumulative frequency of log2 normalized gRNA read counts from six CRISPR/Cas9 knockout screens of distinct ATRT cell lines and the corresponding plasmid reference. (D) Bar graph showing the total number of identified dependencies in six ATRT cell lines at indicated false discovery rate (FDR). (E) Volcano blot illustrating depletion and enrichment on the gene level averaged across CRISPR/Cas9 screens in six ATRT cell lines. Known non-essential and pan-essential genes, as well as genes that are not known to be pan-essential but are significantly depleted across ATRT cell lines (≥ 4 cell lines) are color-coded. (F) Top 10 most significantly enriched gene ontology terms in ATRT-specific genetic dependencies. (G) Drug Gene Interaction database categorization of ATRT-specific essential genes into druggable categories, with 4 sub-categories depicted. For each category, selected genes are listed. See also Figures S1, and S2; Tables S1, and S2.

To identify potential drug targets for ATRTs in an unbiased manner, we next performed genome-wide CRISPR/Cas9 knockout screens using the previously described Brunello library (Doench et al., 2016) in order to derive a list of context-specific genetic dependencies in ATRTs (Figure 1B). Each cell line was transduced with the library at a MOI of 0.3 and cultured for 21 days in three technical replicates. Massive parallel sequencing was subsequently used to quantify gRNA integrants. Replicates agreed well for six out of seven screens, indicating consistent screen results in all cell lines except CHLA02 (Pearson r < 0.7 for technical replicates; Figure S2A; Table S2). The CHLA02 line was therefore omitted from further screen analyses. As expected, deep sequencing revealed a significant reduction in the diversity of gRNAs in ATRT screens as compared to the initial Brunello library plasmid pool from which the library virus was produced (Kolmogorov-Smirnov test *p* < 0.0001 for all screens, Figure 1C; Figure S2B). Known common essential genes showed strong gRNA depletion versus negative controls, indicating good screen signal for gRNAs that affect cell proliferation. On the gene level, we identified an average of 1,055 (range 480-1529) essential genes at FDR of 10 % across ATRT cell lines (Figure 1D; Table S2), showing a significant enrichment for pan-essential genes involved in key cellular processes (Figure S2C, D). After correction for pan-essential genes, we identified 124-405 context-specific essential genes (671 genes in total) per ATRT cell line, including 197 essentials that were depleted in four or more cell lines (Figure 1E; Figure S2E). Further gene ontology analyses of these essentials revealed significant enrichment of distinct functional annotations including dependencies associated with chromatin remodeling, being well in line with the notion that ATRTs are an epigenetically driven disease (Figure 1F; Figure S2F,G). We next aimed to nominate potential targets for pharmacologic inhibition in ATRTs. To first test the coherence of genetic and chemical vulnerabilities in our dataset, we used the Drug Signatures Database (Yoo et al., 2015) to benchmark our screens based on the previously described sensitivity of ATRTs to tyrosine kinase inhibitors (Oberlick et al., 2019; Torchia et al., 2016). Using a set of non-ATRT cell lines previously screened with the Brunello library as a reference (DeWeirdt et al., 2020), we found 16 kinase inhibitor-induced signatures such as that produced by the multi-kinase inhibitor lenvatinib to be significantly enriched (nominal *p* value < 0.1) in ATRT-specific depleted genes (Figure S2H). Reassured by the concordance of our screens with previous results, we next searched the druggability of 671 context-specific essential genes from ATRT cell lines using the Drug Gene Interaction database (DGIdb) (Cotto et al., 2018), and found that 245 of these genes are in druggable categories (Figure 1G). Among these are 37 genes which can be modulated with clinically available compounds, including cyclin-dependent kinases, receptor tyrosine kinases (RTKs), and regulators of p53 signaling. Taken together, our functional genomics approach provided the basis for subsequent functionally-instructed investigations of molecular-based targeted therapies in ATRTs.

### Gene expression is the primary predictor for genetic dependencies in ATRTs

The ability to predict gene essentiality from tumor features can provide further insights into the molecular biology of a given tumor and guide patient stratification. Best predictors for essentialities in cancer cell lines are genetic drivers (oncogene addiction), synthetic lethal interactions, and most notably mRNA expression levels that account for the vast majority of genetic dependencies (McDonald et al., 2017; Tsherniak et al., 2017). We therefore first used an *in silico* analysis to predict oncogenic mutations present in our ATRT cell lines, yielding a list of 30 potentially tumor-driving genetic alterations (Table S1). Only one of those mutations coincided with a significant gene dependency, which was, however, also present in three other cell lines not carrying this mutation. Thus, oncogene addiction is not a predictor of gene essentiality in our cell lines, being in line with the notion that loss of *SMARCB1* is the only recurrent tumor-driving event in ATRTs.

In contrast, we identified several previously described synthetic lethal dependencies (McDonald et al., 2017). All *TP53* wild-type ATRT cell lines were sensitive to loss of negative regulators of p53 signaling *MDM2* and *MDM4* (Figure 2A), representing a vertical pathway synthetic lethal dependency. Of note, loss of *TP53* enhanced proliferation of all ATRT cell lines as the top enriched gene across our screens, further supporting previous results describing the p53 axis as a therapeutic vulnerability in rhabdoid tumors (Howard et al., 2019). As an example for an expression driven paralog lethality, we identified low expression of *CDK6* to predict *CDK4* dependency in ATRT cells (Figure 2B), a feature also seen in other tumor types regardless of tissue origin (McDonald et al., 2017). Due to the overall low mutational burden and stable genomes of ATRTs, we reasoned that the majority of dependencies might be driven by high expression of the corresponding gene, as is true for most cancer dependencies (McDonald et al., 2017). Indeed, we observed an overall negative correlation between DNA promoter methylation and mRNA gene expression across ATRT cell lines (Pearson’s *r* = −0.38). We next correlated mRNA expression with gene level dependency across all ATRT cell lines, revealing a negative correlation, where high gene expression is associated with gene essentiality (Figure 2C). These data suggest that dependencies shared across all ATRT lines are best explained by high expression of the corresponding gene, but it does not explain whether gene expression differences between ATRT cell lines also account for distinct gene essentiality in subsets of ATRT lines. We therefore correlated dependencies with mRNA expression for 671 ATRT-specific essential genes by determining the Pearson correlation coefficient for each gene individually across ATRT cell lines (Figure 2D). We observed a statistically significant shift towards a negative correlation of gene dependency and expression, suggesting that essentiality may be predicted to a useful extent by high mRNA expression for some genes. For several clinically targetable genes, such as *CCND2, FGFR2*, and *ERBB4* (Figure 2E), the correlation is strong across the ATRT cell lines, suggesting that expression of these genes may be useful as markers for patient stratification.

**Figure 2.**
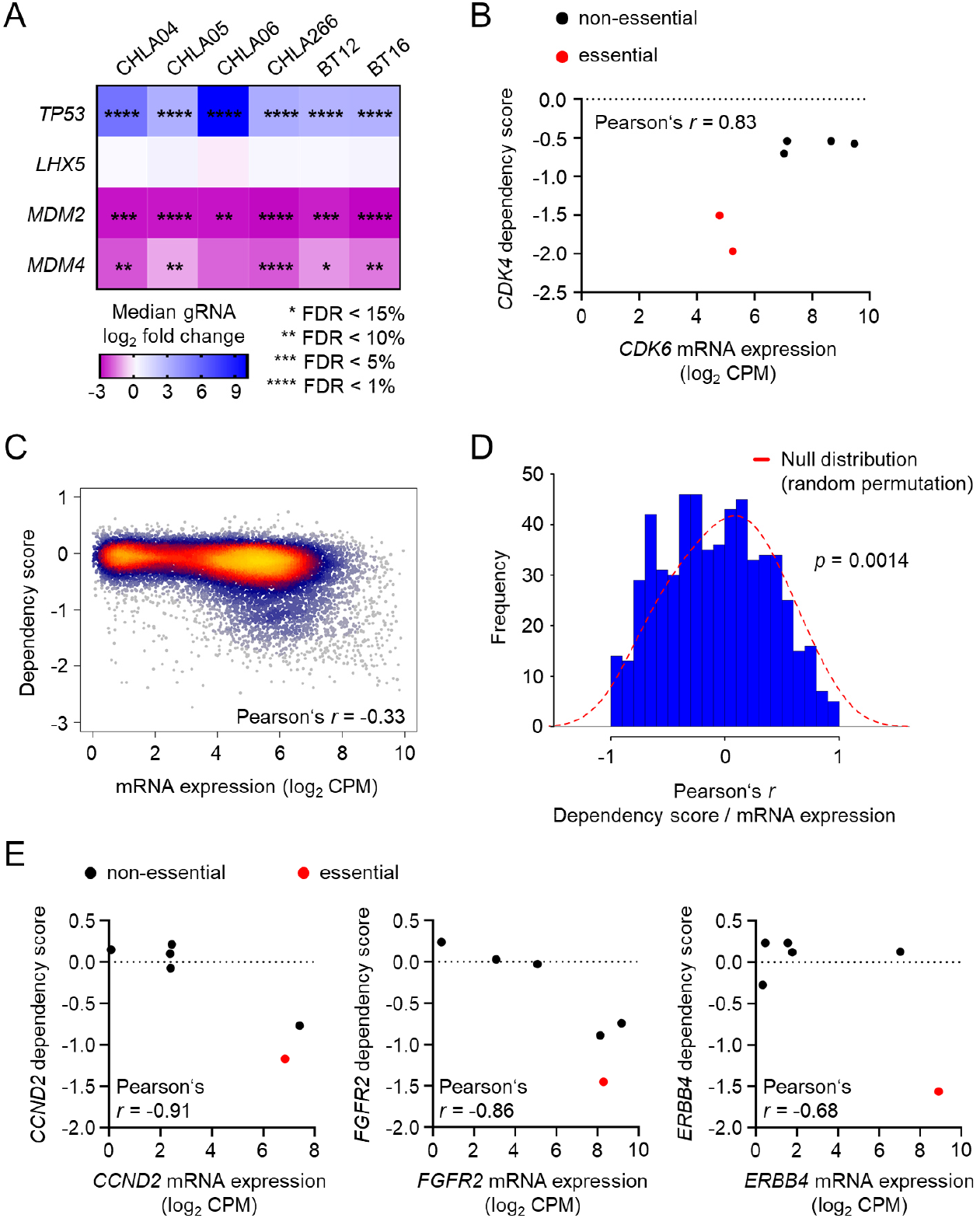
Gene expression is the primary predictor for genetic dependencies in ATRTs. (A) Heatmap illustrating the enrichment of *TP53* and the depletion of negative regulators of p53 signaling *MDM2* and *MDM4* in CRISPR/Cas9 knockout screens of ATRT cell lines. (B) *CDK4* dependency scores plotted versus mRNA expression of *CDK6* in ATRT cell lines. (C) Scatterplot based on a two-dimensional Kernel density estimation illustrating the correlation of dependency scores and mRNA expression averaged across six ATRT tumor cell lines. (D) Correlation of dependency scores and mRNA expression for 671 ATRT-specific genes. The *y* axis represents the number of genes, and the *x* axis illustrates the corresponding Pearson correlation coefficients in 0.1 bins. The dotted line shows the empirical null distribution. Significance was determined using the Wilcoxon rank sum test with continuity correction. (E) Selected genes with strong expression-based dependency profiles across ATRT cell lines. Genes determined to be essential by the STARS algorithm are indicated in red.

### CRISPR screens guide the identification of chemical dependencies in ATRTs

We reasoned that the list of ATRT-specific genetic dependencies could be used to select potential drugs for a molecular-based therapy in ATRTs. Using the DGIdb, we therefore generated a chemical library of 44 distinct small-molecule inhibitors, consisting of 37 compounds described as inhibitors of context-specific genetic dependencies in ATRT cell lines, five compounds previously shown to be efficacious in ATRTs in a subgroup-dependent manner (Torchia et al., 2016), and two broadly cytotoxic agents (Figure 3A). Using a three dose drug screen, we determined the effects of this library on the viability of seven ATRT cell lines and 12 non-ATRT control cell lines representing a diverse set of brain, blood, lung, and soft tissue tumors (Figure S3). To account for substantial differences in the division rate of tested cell models (doubling time range 23 h – 135 h for all cell lines; ATRT-SHH: 113 h ± 24; ATRT-TYR/MYC: 57 h ± 19), we used growth rate inhibition (GR) and area-over-the-curve-response curves (AOC) metrics previously shown to be independent of cell division rate (Hafner et al., 2016) to measure potency and efficacy of each small molecule. As evidenced by unsupervised hierarchical clustering, the majority of ATRT cell lines showed a highly homogenous response to our drug library, and inhibitors with overlapping target profiles clustered together, suggesting on-target activity (Figure 3B; Table S3). Overall, all inhibitors that were picked on the basis of ATRT CRISPR screens (n = 37) were more potent in ATRT cell lines as compared to non-ATRT cell lines (Wilcoxon rank sum test *P* = 2.2 × 10^−16^; Figure 3C), providing evidence for a tumor-specific sensitivity to these classes of compounds.

**Figure 3.**
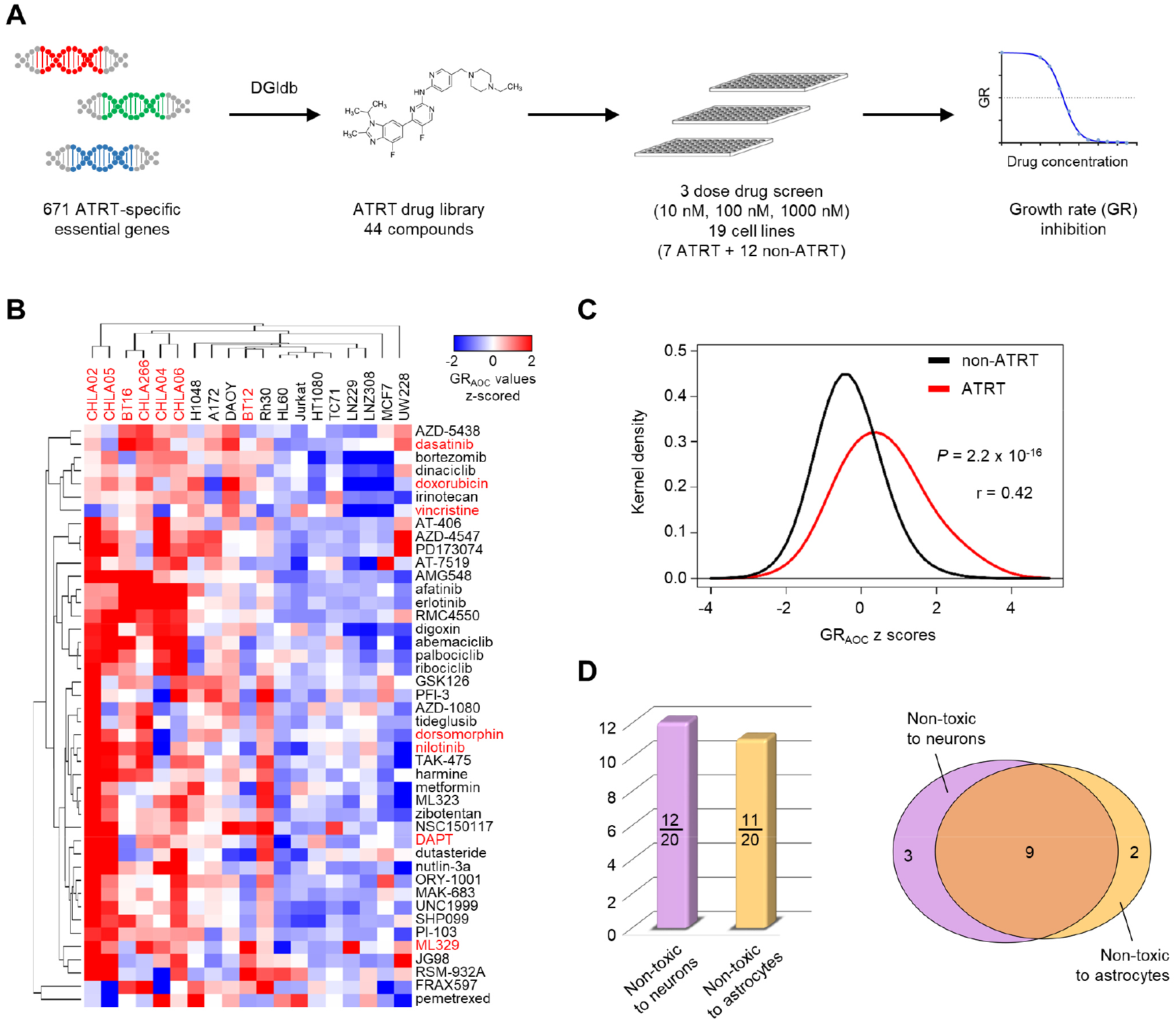
CRISPR screens guide the identification of chemical dependencies in ATRTs. (A) Schematic illustrating the identification of an informed drug library based on ATRT-specific genetic dependencies and downstream drug screening. (B) Unsupervised hierarchical clustering (1 minus Pearson correlation, average linkage) of z-scored growth rate inhibition area-over-the-curve values (GR_AOC_) derived from a three dose drug screen in 19 different cancer cell lines. ATRT tumor cell lines are indicated in red. Small-molecules chosen based on CRISPR/Cas9 screening results (n=37) are shown in black. (C) Kernel density estimation of z-scored GR_AOC_ values from CRISPR/Cas9 screen-based small-molecules (n=37) grouped by ATRT and non-ATRT cell lines. Significance was determined using the Wilcoxon rank sum test with continuity correction (r = Wilcoxon effect size). (D) Neurotoxicity testing of selected small-molecules (GR_max_ < 0.5 in at least four ATRT cell lines, n=20) on postmitotic cerebellar granule neurons and human astrocytes. Bar graph on the right shows the number of compounds that do not significantly inhibit viability of either granule neurons or astrocytes as determined by one-way ANOVA compared to DMSO control (n=4, duplicate for each condition). Venn diagram on the right shows the overlap of small-molecules that are not toxic to either neurons or astrocytes. See also Figure S3; Table S3.

In fact, 11 of 37 inhibitors displayed a significantly higher potency in ATRT cell lines compared to non-ATRT cell lines as judged by GR_AOC_ values (*p* < 0.05, two-way ANOVA with Sidak correction). By ranking both on potency across ATRT cell lines and fold change sensitivity over non-ATRT cell lines, the top 10 hits included a diverse set of inhibitors of cyclin-dependent kinases (CDKs), epidermal growth factor (EGF) and phosphoinositide 3-kinase (PI3K) signaling, mitogen-activated protein kinases (MAPK), as well as other inhibitors targeting distinct cellular components such as the Na/K-ATPase and the proteasome.

Many of those drugs significantly inhibit the growth of ATRT cells, but they might also exhibit substantial toxicity to normal cells, thereby limiting their use in a clinical setting. To test the neurotoxicity profile of the most promising compounds in our library, we tested drugs with the most reliable efficacy in GR inhibition across ATRT cells lines (GR_MAX_ < 0.5 in at least 4 ATRT lines, n = 20) on post-mitotic cerebellar granule neurons and astrocytes. Overall, 9 compounds including inhibitors of CDK4/6, MAPK, and EGF signaling did not significantly inhibit viability of normal cells (one-way ANOVA, compared to DMSO control; Figure 3D), suggesting that these classes of compounds represent chemical vulnerabilities of ATRTs with a broad therapeutic window.

ATRT subgroups exhibit distinct chemical sensitivities (Torchia et al., 2016). We therefore re-analyzed both our genetic and chemical dependency screens, focusing on differential vulnerabilities of ATRT-SHH (group1) and ATRT-TYR/MYC (group 2). We first used multi-set intersection analyses (Wang et al., 2015) to investigate a potential subgroup-specific, differential enrichment of context-specific genetic dependencies from our ATRT CRISPR screens. All possible intersections of essentialities between any two ATRT cell lines were highly significant (Bonferroni adjusted *p*-values < 2.39 × 10^−101^, fold enrichments > 33.67; Figure 4A), possibly underlining the common oncogenic event shared by all ATRTs. However, we did not find any apparent clustering of dependency intersections associated with ATRT subgroups. For example, while ATRT-SHH cell lines (CHLA04 and CHLA05) share 152 genetic dependencies, CHLA05 ATRT cells share many more essentialities with most of ATRT-TYR/MYC cell lines (average 173, range 72 – 225), and CHLA04 cells show a similar overlap with these cell lines (average 131, range 76 – 157). In line with these findings, pairwise dependency set similarities as measured by Jaccard indices and corresponding *p*-values of intersections do not support a subgroup-driven enrichment of single gene dependencies in ATRTs (Figure 4B). However, chemical vulnerabilities, especially those involving multi-targeted inhibitors, might actually not be directly related to single gene dependencies. Analyzing the mean GR_AOC_ in our 3 dose drug screen testing 44 small-molecules, there was a highly significant positive correlation between ATRT-SHH and ATRT-TYR/MYC cell lines (Figure 4C), indicating that cell models from distinct ATRT subgroups show similar sensitivity to most drugs in our library. To further strengthen these results, we performed detailed 15 point GR dose-response analyses for selected compounds from our library that might show selectivity for ATRT subgroups based on preliminary data from our 3 dose drug screen (Figure 4D; Table S3). Both dasatinib and nilotinib, multi-targeted kinase inhibitors previously described to selectively inhibit viability of ATRT-TYR/MYC cell lines (Torchia et al., 2016), did not show any significant difference in their ability to inhibit the growth rate of ATRT-SHH and ATRT-TYR/MYC cell lines (unpaired t test of GR_50_ values). In essence, none of the selected drugs from our informed chemical library including inhibitors of RTKs and CDKs displayed a significant selectivity for a specific ATRT subgroup. Together, these data strongly suggest that chemical vulnerabilities are a result of entity-specific molecular mechanisms in ATRTs, rather than a consequence of epigenetic and transcriptional features defining ATRT subgroups.

**Figure 4.**
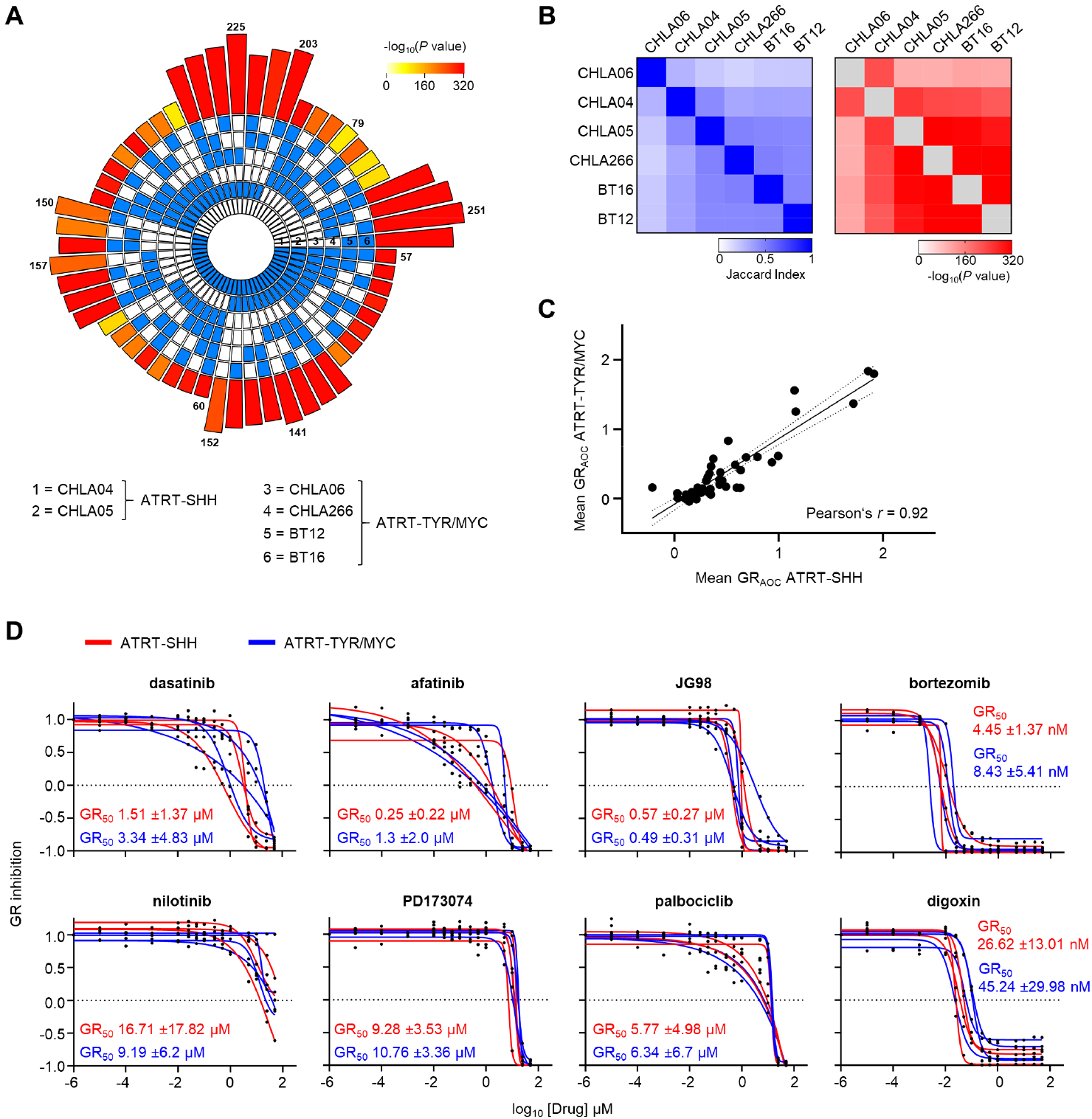
Chemical vulnerabilities are independent from molecular subgroups in ATRTs. (A) Multi-layer circular plot illustrating all possible intersections of ATRT-specific essential genes from six ATRT cell lines. The inner panels show absence (white) or presence (blue) of a specific cell line in a given intersection. The outer bar height represents the intersection size, and the bar color intensity represent statistical significance. (B) Heatmaps illustrating the Jaccard indices (left) and the corresponding significance (right) of pairwise intersections of ATRT-specific essential genes. The order of the heatmap was determined by unsupervised hierarchical clustering (1 minus Pearson correlation, average linkage) of the samples based on their Jaccard indices. (C) Scatterplot showing the correlation of mean GR_AOC_ values from ATRT-SHH and ATRT-TYR/MYC cell lines for all 44 small-molecules from the ATRT drug library tested in a 3 dose drug screen (see Figure 3). (D) 15 point growth rate dose-response-curve analyses for selected small-molecules in ATRT-SHH (CHLA02, CHLA04, CHLA05) and ATRT-TYR/MYC (BT12, BT16, CHLA06, CHLA266) cell lines. Mean GR50 values for ATRT-SHH and ATRT-TYR/MYC subgroup cell lines are shown.

### CDK4 and CDK6 are distinct predictors of CDK4/6 inhibitor sensitivity in ATRTs

CDK4/6 inhibitors were among the most potent small-molecules in our drug screen, which preferentially inhibited growth of ATRT cell lines, while sparing normal neural cell types. CDK4/6 inhibition is predicted to lead to cell cycle arrest in G1 phase, and this is also true for ATRT cell lines (Figure S4A). To first test the general feasibility of CDK4/6 inhibition as a therapeutic strategy for ATRTs, we again performed detailed dose-response analyses on several ATRT cell lines including ATRT-311FHTC, a PDX-derived ATRT-SHH cell line (Brabetz et al., 2018), using abemaciclib, the most potent CDK4/6 inhibitor in our drug screen. In keeping with our above findings using palbociclib (Figure 4D), abemaciclib exerted a cytotoxic effect in all cell lines, but did not show a significant difference in its potency towards ATRT-SHH and ATRT-TYR/MYC cells (Figure 5A). Furthermore, ATRT cell lines showed a similar sensitivity to abemaciclib in colony formation assays as MCF7 cells (Figure 5B), an ER^+^ breast cancer line known to be highly susceptible to CDK4/6 inhibition (Finn et al., 2009). To test the efficacy of abemaciclib in inhibiting ATRT tumor growth *in vivo*, we next employed a orthotopic xenograft mouse model using BT16 as a model for ATRT-MYC. Mice treated orally with abemaciclib (75mg/kg) for 3 weeks had significantly prolonged survival compared with corresponding vehicle-treated controls (*p* = 0.0202; Figure 5C), arguing that abemaciclib is able to cross the blood-brain-barrier as suggested previously (Raub et al., 2015) to inhibit ATRT tumor growth *in vivo*.

**Figure 5.**
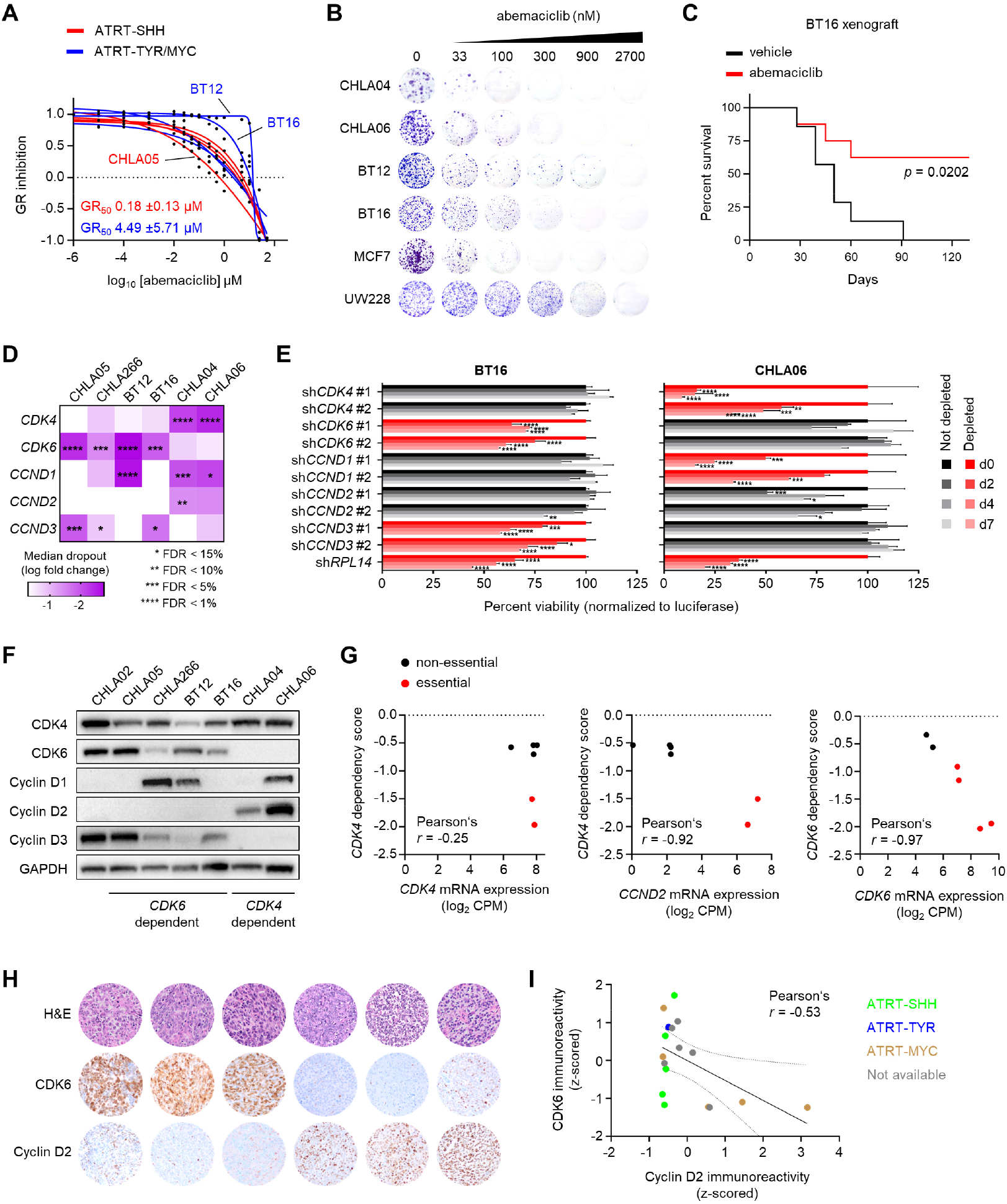
CDK4 and CDK6 are distinct predictors for CDK4/6 inhibitor sensitivity in ATRTs. (A) 15 point growth rate dose-response-curve analyses for the CDK4/6 inhibitor abemaciclib in ATRT-SHH (CHLA02, CHLA04, CHLA05, ATRT-311FHTC) and ATRT-TYR/MYC (BT12, BT16, CHLA06, CHLA266) cell lines. (B) Colony-formation assay of several ATRT cell lines treated with abemaciclib. An ER^+^ breast cancer cell line (MCF7) and a medulloblastoma cell line (UW228) are also shown as reference. (C) Kaplan-Meier survival analysis of a BT16 cell-based intracranial transplantation mouse model treated daily with 75mg/kg abemaciclib (n = 8) or vehicle (n = 7), monitored for 130 days after tumor cell transplantation. (D) Heatmap illustrating gene level log fold changes and corresponding FDR statistics for CDK4, CDK6, and all D type cyclins in ATRT CRISPR knockout screens. (E) Bar graphs showing the effect of shRNA-mediated knockdown of *CDK4, CDK6, CCND1, CCND2*, and *CCND3* in BT16 and CHLA06 cells (n = 3 independent experiments; 2 distinct shRNAs for each target; Two-way ANOVA with Dunnett correction for multiple comparisons). As a control, a shRNA for knockdown of the pan-essential gene *RPL14* was included, and values were normalized to the effect of a shRNA targeting the luciferase gene. Genes in red were determined to be essential in the corresponding cell line by CRISPR screen. Data are mean ±SEM. (F) Western blot analyses showing the protein expression levels of CDK4, CDK6, and D type cyclins in the indicated ATRT cell lines. (G) Correlation analyses to illustrate *CDK4* or *CDK6* dependency prediction by *CCND2* and *CDK6* mRNA expression. Pearson’s r values are shown. (H) Representative H&E stains and immunohistochemistry for CDK6 and cyclin D2 (DAB as brown chromogen) in sections from FFPE ATRT tissue from 6 distinct patients ordered by column. (I) Correlation analysis of immunoreactivity for CDK6 and cyclin D2 in primary ATRT tissue from a total of 17 patients. See also Figure S4.

To gain further insights into the mechanisms underlying CDK4/6 inhibitor sensitivity in ATRTs, we first re-investigated our CRISPR screen dataset. All ATRT cell lines depend on either *CDK4* or *CDK6* expression in a mutually exclusive manner, a pattern recapitulated by most human cancer cell lines (Figure S4B), and this did not correlate with the three molecular subgroups (Figure 5D). Validating this by shRNA-based knockdown approaches, we found that knockdown of either *CDK4* or *CDK6* differentially inhibited the proliferation of distinct ATRT cell lines in line with our CRISPR screen results (Figure 5E). In addition, BT16 and CHLA06 cell lines were also differentially dependent on distinct D type cyclin paralogs, further supporting a heterogeneous mechanism underlying CDK4/6 inhibitor sensitivity in *CDK4*-versus *CDK6*-dependent ATRTs. We reasoned that differential expression of members of the CDK/D-type cyclin axis might be responsible for this heterogeneity, as already suggested by our findings that low expression of *CDK6* predicts *CDK4* dependency (Figure 2B). We validated this synthetic lethal interaction by investigating protein expression levels in ATRT cell lines, which also uncovered an unexpected heterogeneity in the expression of CDKs and D-type cyclins (Figure 5F). Indeed, we identified cyclin D2 and CDK6 expression as a molecular predictor for *CDK4*- and *CDK6*-dependent cell lines: while *CDK4* expression does not predict *CDK4* essentiality, *CDK4*-dependency strongly correlated with *CCND2* expression (Figure 5G) and a complete lack of CDK6 protein detection. In contrast, *CDK6*-dependent ATRT cells show high expression of CDK6, both on the mRNA and protein level. Notably, this correlation is conserved across human cancer cell lines, where low expression of *CDK6* is correlated with *CDK4* dependency, whereas high expression of *CDK6* is a predictor for *CDK6* dependency itself (Figure S4C). However, *CDK4* dependency across diverse lineages of tumor cell lines is clearly correlated with high expression of cyclin D1 rather than cyclin D2, contrasting our results from ATRT cell lines. To further validate our surprising data in clinical samples, we tested protein expression of CDK6 and cyclin D2 in 17 primary ATRTs histology samples using immunohistochemistry (Figure 5H). In fact, we found a clear negative correlation of CDK6 and cyclin D2 protein expression (Pearson’s r = −0.53), where high CDK6 expression correlated with low cyclin D2 expression, and vice versa (Figure 5I). Together, our data suggest that ATRT cells segregate into two groups with distinct dependencies within the CDK4/6-D type cyclin axis, making them highly susceptible to CDK4/6 inhibition.

### CDK4 and CDK6 dependency drives heterogeneity in response to CDK4/6 inhibition

Finally, we aimed to investigate to what extent these differences in CDK4/6 dependency affect downstream molecular mechanisms in ATRT cells, and, from a clinical perspective, whether this will predict the response of ATRT to CDK4/6 inhibition. Grouping ATRT cell lines according to *CDK4* or *CDK6* dependency, we found a highly significant difference in global gene expression profiles with a total of 2,643 dysregulated genes (│log_2_ fold change│ ≥ 1; adjusted *p* < 0.01), which we did not detect by randomly assigning *CDK4* or *CDK6* dependency status to ATRT cells (Figure 6A, Table S4). While most enriched gene ontologies were clearly associated with the regulation of signaling pathways, mutually exclusive enrichment of functional terms such as MAPK or G protein-coupled receptor signaling further supported distinct biological processes active in *CDK4*- and *CDK6*-dependent ATRT cells (Figure 6B, Table S4). We further assessed possible cellular heterogeneity in the response to CDK4/6 inhibition related to *CDK4* and *CDK6* dependency. To this end, we performed single cell RNA-seq (scRNA-seq) on *CDK4*- and *CDK6*-depencent ATRT cells (BT12, BT16, CHLA06) treated with abemaciclib or vehicle. Data from an average of 4,383 DMSO-treated and 3,816 abemaciclib-treated cells per cell line that met quality control criteria were used for transcriptomic analysis. Visualization of the data from vehicle-treated cells via uniform manifold approximation and projection (UMAP) revealed that ATRT cells were broadly segregated according to their *CDK4* and *CDK6* dependency status (Figure 6C). Distinct cell populations recapitulated gene expression profiles with respect to marker genes of distinct CDK dependencies (Figure 6D, Figure S5A). Further analyses identified nine transcriptionally distinct cell cluster that were unrelated to cell cycle phase (Figure 6E, Figure S5B). Of note, clusters 1, 4, 5, and 6 contained cells from all ATRT cell lines, but were largely dominated by *CDK6*-dependent BT12 (cluster 1, 6) and BT16 cells (cluster 1, 4, 5), whereas cluster 8 only contained cells from BT12 cells. In contrast, transcriptionally distinct cell clusters 2, 3, 7, and 9 were exclusively comprised of CHLA06 cells, thus containing the vast majority of this ATRT cell line. These results suggest that ATRT cell lines present inter- and intratumoral heterogeneity that is at least in part driven by differential *CDK4* and *CDK6* dependency.

**Figure 6.**
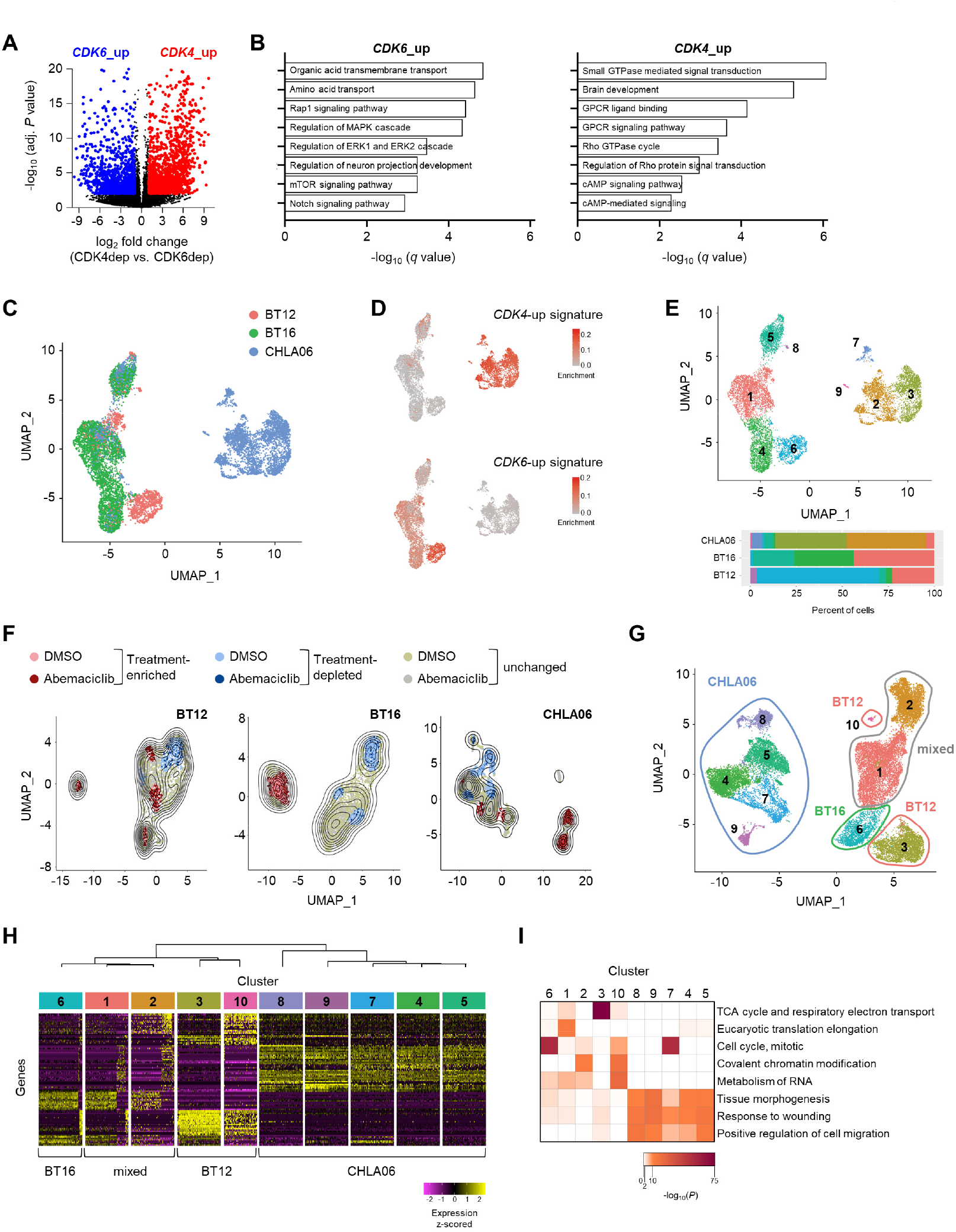
CDK4 and CDK6 dependency drives heterogeneity in response to CDK4/6 inhibition. (A) Volcano plot illustrating global gene expression differences in *CDK4*- and *CDK6*-dependent ATRT cell lines. (B) Selected gene ontology terms enriched in *CDK4*- and *CDK6*-dependent ATRT cell lines. (C) UMAP dimensionality reduction of scRNA-seq data from control BT12, BT16, and CHLA06 ATRT cell lines. (D) UMAP embedding of scRNA-seq data from control BT12, BT16, and CHLA06 cell lines with enrichment projection of gene signatures indicative of *CDK4* or *CDK6* dependency derived from bulk RNA-seq data (genes with log_2_ fold change > 4). (E) UMAP projection of single cell profiles from control BT12, BT16, and CHLA06 cell lines clustered by transcriptional similarities (top). Bar chart indicates cell cluster composition of cell lines (bottom). (F) UMAP embedding with superimposed Kernel density estimation for abemacilib- and vehicle-treated cells individually plotted for each cell line. The density estimate for each cell from treated and untreated conditions was calculated, and a Δ density contrast (abemaciclib – vehicle) was constructed to determine cell populations either enriched (> 80^th^ percentile of all Δ values) or depleted (< 20^th^ percentile) under abemaciclib treatment. (G) UMAP embedding and clustering of scRNA-seq data from BT12, BT16, and CHLA06 cells treated with vehicle (DMSO) or abemaciclib. Dominating cell type for clusters is indicated. (H) Heatmap showing unsupervised hierarchical clustering of cell cluster and the expression of the top 20 markers per cluster across a total of 24,597 single cells from BT12, BT16, and CHLA06 cells treated with vehicle (DMSO) or abemaciclib. (I) Heatmap illustrating enrichment of top 10 process networks in marker genes for each cell cluster.

We next incorporated data from the same ATRT cell lines treated with abemaciclib and found that, in each cell line, abemaciclib treatment induced a strong shift of discrete subpopulations that were either enriched or depleted (Figure 6F), suggesting that intratumoral heterogeneity contributes to differential response to CDK4/6 inhibition. In addition, using all cell lines together, cells still clustered by cell line rather than treatment condition (Figure S5C), suggesting that CDK4/6 inhibition does not impose major transcriptional changes in ATRT cells. However, abemaciclib treatment induced a strong G1 cell cycle arrest in all cell lines (Figure S5D, E). Clustering based on transcriptional features revealed 10 distinct cell clusters, still largely segregated by differential dependency on *CDK4* or *CDK6* (Figure 6G). Similar to vehicle-treated ATRT cell lines, most clusters were dominated by one cell line. However, we did not observe a cluster-specific association with a particular treatment condition (Figure S5F), again suggesting that cluster identity is mainly driven by cell intrinsic mechanisms rather than CDK4/6 inhibition. A notable exception is cluster 7, and to lesser extent cluster 6, which were greatly diminished under CDK4/6 inhibition. This is in line with the fact that these clusters were almost exclusively comprised of cells in S or G2/M phase (Figure S5G), which are expected to be preferentially depleted under CDK4/6 inhibition. Cell clusters dominated by the same cell line were highly similar with respect to their marker gene expression (Figure 6H). Accordingly, we observed distinct molecular phenotypes enriched in clusters dominated by different cell lines, including mitochondrial energy processes, RNA metabolism, and cell migration (Figure 6I). These data indicate that, apart from G1 cell cycle arrest, ATRT cells respond differently to CDK4/6 inhibition, and that differences in response might at least in part be attributed to gene expression profiles associated with *CDK4* and *CDK6* dependency in ATRTs.

## Discussion

High-throughput next-generation sequencing is a powerful approach to discover molecular alterations in disease entities and to drive translational innovations, e.g. by molecular-based patient stratification and/or molecular-driven therapies. Yet, the translational impact of next-generation sequencing can only be fully exploited if actionable genetic alterations are present in the investigated tissue. Consequently, the impact on molecular-driven clinical innovations in diseases without actionable genetic alterations, like ATRT, is rather limited. Furthermore, genomics or transcriptomics alone provide a “static” molecular analysis and thereby leaves out any alterations by adaptivity. Functional genomics and subsequent validation steps in clinically relevant models might bridge these limitations. We applied this novel approach to ATRTs, an incurable pediatric brain tumor. Our study provides an unprecedented view of genetic dependencies in ATRTs at genome scale, and illustrates various molecular mechanisms underlying tumor cell-specific dependencies.

We identified known molecular feature/dependency interactions (McDonald et al., 2017; Tsherniak et al., 2017) in the ATRT cell lines, confirming their utility for the purpose of our study. The genetic profile combined with our CRISPR screen data (Figure S1, Figure 2) suggests inactivation of *SMARCB1* as the sole recurrent genetic driver in ATRTs supporting previous studies (Johann et al., 2016; Torchia et al., 2016). In contrast, synthetic lethal interactions, in which a non-self-association is correlated with a specific genetic dependency, account for some of the observed essentialities. All ATRT cell lines were highly sensitive to alterations in the p53 signaling pathway (Figure 2), in line with observations that rhabdoid tumors are particularly sensitive to MDM2/4 inhibition (Howard et al., 2019). Furthermore, the CDK4/6 signaling axis in ATRTs reflects distinct mechanisms of gene essentiality, contributing to mutual exclusive dependencies on either *CDK4* or *CDK6*. While *CDK4* dependency is strongly correlated with low *CDK6* expression, representing a synthetic lethal interaction present in diverse sets of tumor entities (McDonald et al., 2017), *CDK6* essentiality itself is an mRNA expression-based genetic dependency. Small-molecule inhibitor screening revealed a striking sensitivity of ATRT cells over non-ATRT cells to compounds that we selected based on context-specific dependencies from ATRT CRISPR screens (Figure 3). We are aware that genetic dependencies such as *CDK4* and *CDK6* are not ATRT-specific *per se* and occur in a variety of distinct tumor entities. However, the unique combination of context-specific dependencies might well be specific for ATRT cells as suggested by our drug screen. In particular, the potential mechanisms of escape for ATRT from targeting of any of these dependencies may share some commonalities in ATRT versus other cancers. Thus, our data might pave the way for novel investigations on resistance mechanisms and potential synergistic combinations in ATRTs.

Of note, the dependencies identified by CRISPR screening and corresponding chemical vulnerabilities in our study do not correlate with specific molecular ATRT subgroups (Figure 4). This finding is in contrast to a previous study observing subgroup-specific sensitivities in ATRTs. Yet, this study relied on relative viability assays (Torchia et al., 2016). As the quantification of drug assays using relative cell counts can be largely confounded by the cell division rate of *in vitro* model systems (Hafner et al., 2016; Hafner et al., 2017), we suggest that the lack of subgroup-specific chemical vulnerabilities in our study are due to our different experimental approach. The impact of cell division rates on quantifications of drug sensitivities might be of particular importance when using ATRT cell lines, because these cells present a wide range of doubling times, with ATRT-SHH cell lines dividing substantially slower than ATRT-TYR/MYC cells, thereby preventing reliable quantification of drug potencies based on relative cell counts. Our analyses using growth rate inhibition drug dose-responses (Figure 3) are in line with intersection analyses of genetic dependencies from ATRT cells, which did not provide any evidence for subgroup-specific enrichment. Since the best common genetic predictor for our identified subgroup-independent ATRT dependencies is loss of the tumor suppressor *SMARCB1*, we suggest to investigate these vulnerabilities in other *SMARCB1*-deficient tumors beyond ATRTs, e.g. extracranial rhabdoid tumors, epithelioid sarcomas, or malignant peripheral nerve sheath tumors (Margol and Judkins, 2014).

According to our data, CDK4/6 inhibitors are among the most promising candidates for molecular-driven therapies in ATRT patients as these compounds showed preferential efficacy in ATRT cells over non-ATRT cells, while sparing normal brain cell types. In addition, CDK4/6 inhibitors have a well-known safety profile as they have approval for treatment of certain types of breast cancer (Wu et al., 2020) and cross the blood-brain barrier (Nguyen et al., 2019).

Our study reveals for the first time a mutual exclusive dependency pattern of *CDK4* and *CDK6* in ATRT cells (Figures 5), which correlates with a distinct downstream gene expression profile (Figure 6). Since all currently available CDK4/6 inhibitors have similar affinities to both CDK4 and CDK6 (Marra and Curigliano, 2019), it is unlikely that the potency of a current inhibitor is influenced by *CDK4* and *CDK6* dependency in ATRT cells, and we detected no such tendency in ATRT cell lines. However, we identified distinct bulk and single cell gene expression profiles corresponding to distinct CDK dependencies in ATRT cell lines, and primary ATRTs might display these differences as well as evidenced here by negative correlation of CDK6 and cyclin D2 protein expression levels (Figure 5). While clonal heterogeneity has been recently suggested also in primary ATRTs (Jessa et al., 2019), our data show that this type of heterogeneity shapes the molecular response of ATRT cells to CDK4/6 inhibition. This suggests that *CDK4*- and *CDK6*-dependent ATRT cells might display differential mechanisms of synthetic lethal interactions and resistance in response to CDK4/6 inhibition, that in turn might enable potential combination or sequential therapies. Genome-scale perturbation screens will be a promising experimental approach to answer these fundamental questions.

In summary, we present a comprehensive ressource for investigations on molecular-based therapeutic strategies for ATRTs. Our results particularly warrant further clinical translation of CDK4/6 inhibitors as a promising therapeutic approach for ATRTs. Furthermore, this study might serve as a blueprint how target discovery by large-scale genetic screens and their comprehensive validation by independent methods and in clinically relevant samples contribute to the development of functionally-instructed molecular therapies for diseases lacking actionable genetic events.

## Acknowledgements

We thank Sarah Hendel, Heike Pfrommer, and Yeliz Donat for excellent technical assistance. Parts of this study were funded by the Deutsche Forschungsgemeinschaft (ZUK63 to D.J.M), the German Scholars Organisation (GSO/EKFS 5 to G.T.), the Else Kröner-Fresenius Stiftung (Else Kröner-Forschungskolleg Tübingen, 2015_Kolleg_14, to G.T.), the Adolf Leuze Stiftung (to G.T. and D.J.M), the Medical Faculty Tübingen (Demonstratorprojekt Personalisierte Medizin to G.T.), the Deutsche Forschungsgemeinschaft (SCHU 2442/7-1 to U.S.), the Deutsche Krebshilfe (111785 to U.S.), the Wilhelm Sander Stiftung (2018.055.1 to U.S.), and the Fördergemeinschaft Kinderkrebszentrum Hamburg.

## Author Contributions

Conceptualization, D.J.M. and G.T.; Methodology, D.J.M. and G.T.; Formal analysis, D.J.M., S.H., F.T., S.B., J.A., N.C., C.R., M.S., F.P., N.S.P., and M.C.; Investigation, D.J.M., S.H., F.T., B.W., L.A.H.; Resources, J.S., S.S., U.S., D.E.R., and M.T.; Writing – Original Draft, D.J.M. and G.T.; Writing – Review & Editing, D.J.M. and G.T.; Visualization, D.J.M., S.H., and F.T.; Supervision, D.J.M. and G.T.; Project Administration, D.J.M. and G.T.; Funding Acquisition, D.J.M. and G.T.

## Declaration of interests

G.T. reports personal fees (advisory board, speaker’s fees) from AbbVie, Bayer, Bristol-Myers-Squibb, Medac, Novocure, travel grants from Bristol-Myers-Squibb, educational and travel grants from Novocure, research grants from Roche Diagnostics, research and travel grants from Medac. All other authors declare no competing interests. The other authors do not have any conflicts of interest to disclose.

## STAR methods

### RESOURCE AVAILABLILITY

#### Lead contact

Further information and requests for resources and reagents should be directed and will be fulfilled by the Lead Contact, Ghazaleh Tabatabai.

#### Materials Availability

This study did not generate new unique reagents.

#### Data and Code Availability

All datasets generated in this study are either available as supplemental information, or will be made freely available upon acceptance of this manuscript.

### EXPERIMENTAL MODEL AND SUBJECT DETAILS

#### Animals

For orthotopic transplantation of tumor cells, Crl:CD1-Foxn1^Nu/Nu^ nude mice were used (Charles River). All mice were female, and between 8 and 10 weeks old. For treatment procedures, mice were randomly assigned to experimental groups. Sample size calculation was performed by a biometrician. All animal experiments were approved by the Regierungspräsidium Tübingen (N 09/20 G and N 11/20 M). All experiments were conducted in accordance with the animal law. Animals were closely monitored. For reporting, we followed the ARRIVE guidelines (version 2.0).

#### Archived tissue samples from ATRT patients

For immunohistochemistry studies, we used archival tissue from 17 ATRT patients. Both female and male patients were included. Average age of patients was 1.9 years ± 2.5. Molecular subgroups of ATRT tumors had been determined by global DNA methylation profiles within the clinical routine diagnostics using the methylation classifier for central nervous system tumors (www.molecularneuropathology.org). The Ethics Committee of the Medical Faculty of the University Hospital Tübingen approved the study (870/2020A).

#### Cell lines

The following cell lines were used in this study: CHLA02 (RRID: CVCL_B045), CHLA04 (RRID: CVCL_0F38), CHLA05 (RRID: CVCL_AQ41), CHLA06 (RRID: CVCL_AQ42), CHLA266 (RRID: CVCL_M149), BT12 (RRID: CVCL_M155), BT16 (RRID: CVCL_M156), H1048 (RRID: CVCL_1453), A172 (RRID: CVCL_0131), LN229 (RRID: CVCL_0393), LNZ308 (RRID: CVCL_0394), DAOY (RRID: CVCL_1167), UW228 (RRID: CVCL_0572), Rh30 (RRID: CVCL_0041), HL60 (RRID: CVCL_0002), Jurkat (RRID: CVCL_0367), HT1080 (RRID: CVCL_0317), TC71 (RRID: CVCL_2213), MCF7 (RRID: CVCL_0031), and ATRT-311FHTC (Brabetz et al., 2018). CHLA02, CHLA04, CHLA05, and CHLA06 cell were grown in DMEM/F12 media supplemented with B27, EGF, FGF, GlutaMAX and HEPES. CHLA266, BT12, Rh30, and TC71 cells were grown in IMDM media supplemented with 10% FCS and 1x ITS (Insulin-Transferrin-Selenium). BT16, DAOY, HL60, Jurkat, and HT1048 cells were grown in RPMI media supplemented with 10% FCS. A172, LN229, LNZ308, UW228, HT1080, and MCF7 cells were grown in DMEM media supplemented with 10% FCS. All these cell line media were supplemented with gentamycin. PDX-derived ATRT-311FHTC cells were grown in laminin-coated plates in NeuroCult NS-A Basal media supplemented with NS-A Proliferation Supplement, EGF, FGF, Heparin, and penicillin/streptomycin. Cell lines were not authenticated.

## METHOD DETAILS

### DNA methylation profiling by 850k EPIC array

Global DNA methylation from seven human ATRT cell lines was assessed using Illumina Infinium MethylationEPIC 850k arrays according to the manufacturer’s instructions. Raw idat files were read and processed using the *minfi* Bioconductor package in R (3.6.0). Briefly, all probes (n = 865,859) were subjected to quality control metrics, removing all probes with p detection values > 0.01. Furthermore, all probes targeting site on sex chromosomes were removed. Also, all probes with SNPs at the CpG site were removed. In total, 813,439 distinct probes were kept in order to generate β values. For prediction of ATRT subgroups, we first generated several models using machine-learning algorithms using a test cohort of 150 primary ATRTs (Johann et al., 2016). In order to combine 450k and 850k methylation arrays, we generated virtual arrays using the *minfi* package. Employing the 2 % probes (n = 8,513) showing highest variability across all three molecular subgroups in the test cohort, we generated prediction models using the caret package with the following machine-learning algorithms: random forest (rf), stochastic gradient boosting (gbm), support vector machine (SVM), linear discriminant analysis (lda), k-nearest neighbors (knn), nearest shrunken centroids (PAM), decision tree (dt), and classification and regression trees (cart). The three models that predicted ATRT subgroups with highest accuracy in the test cohort (rf, gbm, svm) were used for further prediction studies. Our random forest model was last evaluated on a validation cohort comprising 121 primary ATRTs (Capper et al., 2018). *t*-SNE dimensionality reduction of global DNA methylation data from human ATRT cell lines and a primary CNS tumor reference (n = 2,801) was performed as previously described (Capper et al., 2018).

### Custom oncopanel sequencing

For DNA sequencing, 200 ng of genomic DNA was fragmented to 150-200 bp pairs using ultrasonication on the LE220 Focused-ultrasonicator (Covaris). Library preparation was performed using the SureSelect XT Library Prep Kit (Agilent Technologies) and enrichment of gene of interest was performed using the SureSelect XT Target Enrichment System with custom-designed bait-sets (ssSC v5) covering 708 cancer related genes, 7 promoter regions and selected fusions. The libraries were sequenced as paired-end 75 bp reads on an Illumina NextSeq500 (Illumina) with a sequencing depth of approximately 25 million clusters per sample. DNA raw data QC and processing was performed using the in-house megSAP Pipeline (https://github.com/imgag/megSAP, version 0.1-1223-gf2879e3) combined with ngs-bits package (https://github.com/imgag/ngs-bits, version 2019_08) (Schroeder et al., 2017). Briefly, sequencing reads were aligned to the human reference genome (GRCh37) by BWA-MEM, variants were called using Strelka2 (Kim et al., 2018) and annotated with VEP (McLaren et al., 2016). To obtain high-confidence results filter criteria for variants were defined as a tumor and normal depth of at least 20x, an allelic frequency of 5% or more and a minimum of 3 reads.

### Gene expression profiling using bulk RNA-seq

For RNA sequencing, mRNA fraction was enriched using polyA capture from 200ng of total RNA using the NEBNext Poly(A) mRNA Magnetic Isolation Module (NEB). Next, mRNA libraries were prepared using the NEB Next Ultra II Directional RNA Library Prep Kit for Illumina (NEB) according to the manufacturer’s instructions. The libraries were sequenced as paired-end 50bp reads on an Illumina NovaSeq6000 (Illumina) with a sequencing depth of approximately 25 million clusters per sample. RNA raw data QC and processing was performed using megSAP (version 0.2-135-gd002274) combined with ngs-bits package (version 2019_11-42-gflb98e63). Reads were aligned using STAR v2.7.3a (Dobin et al., 2013) to the GRCh38 and alignment quality was analyzed using ngs-bits. Normalized read counts for all genes were obtained using Subread (v2.0.0) and edgeR (v3.26.6).

### Single cell RNA-seq

The final ATRT cell preparations contained over 78% (ranging from 78 to 88%) live cells at a concentration of 600-1300 cells/μL. Single-cell suspension concentration and cell viability were determined by automatic cell counting (DeNovix CellDrop, DE, USA) using an AO/PI viability assay (DeNovix, DE, USA). scRNA-seq libraries were generated using the 10X Chromium Next gel beads-in-emulsion (GEM) Single Cell 3’ Reagent Kit v3.1 according to the manufacturer’s instructions. Approximately 14,000 cells per sample were loaded on the Chromium Next GEM Chip G, which was subsequently run on the Chromium Controller (10X Genomics, CA, USA) to partition cells into GEMs. Cell lysis and reverse transcription of poly-adenylated mRNA occurred within the GEMs and resulted in cDNA with GEM-specific barcodes and transcript-specific unique molecular identifiers (UMIs). After the breaking of the emulsion, cDNA was amplified using 10 or 11 cycles according to manufacturer’s instructions, enzymatically fragmented, end-repaired, extended with 3′ A-overhangs, and ligated to adapters. P5 and P7 sequences, as well as sample indices (Chromium i7 Multiplex kit, 10X Genomics, CA, USA), were added during the final PCR amplification step. The fragment size of the final libraries was determined using the Bioanalyzer High-Sensitivity DNA Kit (Agilent, CA, USA). Their concentration was determined using the Qubit dsDNA HS Assay Kit (Thermo Fisher Scientific, MA, USA). scRNA libraries were pooled and paired-end-sequenced on the Illumina NovaSeq 6000 platform. Raw data was pre-processed using CellRanger software (10x Genomics). Batch correction was performed using harmony (https://github.com/immunogenomics/harmony). All downstream analyses were performed using the *seurat* package in R (https://github.com/satijalab/seurat).

### Copy number analysis

We followed two distinct strategies in this study to assess copy number changes in ATRT cell lines to complement each other. One, for DNA methylation-based analysis of copy-number variation from 850k EPIC arrays, we used the *conumee* package in R (http://bioconductor.org/packages/conumee/). Two, for somatic copy number alteration detection, we used ClinCNV (version 1.16). By analyzing off-target reads, the tool can generate copy-number information about the whole genome, also for targeted panel sequencing data. A gene with an integer copy-number (CN) of ≥ 4 was defined as amplified. A heterozygous deletion was assumed with an integer CN = 1, a homozygous deletion when CN = 0.

### Gen set and gene ontology analyses

Gene set enrichment analyses (GSEA) were performed using the GSEA software (version 3.0) from the Broad Institute (https://www.gsea-msigdb.org/gsea). All gene sets enriched at FDR < 25 % were considered significant. Single sample GSEA was performed as previously described (Barbie et al., 2009). For gene ontology analyses, we used Metascape (Zhou et al., 2019).

### Genome-wide CRISPR/Cas9 knockout screens

CRISPR/Cas9 screens using a genome-wide knockout gRNA library were performed as previously described (Doench et al., 2016). Briefly, ATRT cell lines were transduced by spinfection with lentiviral particles representing the gRNA library in an all-in-one version containing Cas9 enzyme at a MOI ∼0.3. Cell numbers were estimated to ensure a 500x library coverage in the transduced cell population. On the second day, cells were split to three pseudo-replicates, selected by puromycin for five days, and kept in culture for a total for 21 days to allow for the depletion of cells where gRNA-guided knockouts affect cell survival or proliferation. For each ATRT cell line individually, optimal transduction and growth conditions were pre-determined. Library coverage was kept at 500x at all sub-culturing steps. After 21 days, genomic DNA was isolated using the QIAamp Blood Maxi kit (Qiagen), and subjected to next generation sequencing for quantification of gRNA distribution in the remaining cell population. Illumina sequencing was performed as previously described, and the STARS gene-ranking algorithm was used to determine statistically depleted and enriched gRNAs (Doench et al., 2016).

### Cell culture and growth rate inhibition assays

Cells were cultured in the above-mentioned media under empirically optimized conditions. All cells were cultured in 100 µl supplemented with the corresponding compound at three distinct concentrations (10 nM, 100 nM, 1000 nM) in 96 well plates. Each plate contained a DMSO control that was used to normalize all data from this plate. Each condition on each plate contained 8 replicate values that were averaged upon analysis. We performed the drug screen in two session, and tested the coherence of both sessions constructing a similarity index (Indahl et al., 2018). Cell viability was assessed using CellTiter-Blue (Promega).

### Neurotoxicity assays

For assessment of neurotoxic activity of selected drugs from the ATRT compound library, we used postmitotic cerebellar granule neurons as well as human astrocytes as surrogates for normal cell types of the brain. Granule neurons were generated from wildtype pups at P5. Cerebella were dissected in der the scope, meninges removed, and cerebella were triturated in trypsin/EDTA containing DNase. After washing, cells were counted, and seeded in DMEM/F12 media supplemented with 10 % FCS, 25 mM KCL, 1x GlutaMAX, and Pen/Strep onto polyornithine-coated 96 well plates. On the second day, 10 µM Ara-C were added for 4 days in order to enrich for postmitotic granule neuron cells. In order to test our compounds in a glia-representing lineage, we used normal human astrocytes (NHA) cells from Lonza which were cultured according to manufacturer’s instructions.

### shRNA-based gene knockdown

Two distinct shRNA for each target gene were cloned into pLKO.1 puro (Addgene #8453) according to the corresponding protocol. Lentiviral particles were produced in HEK293T cells, and ATRT cell lines were subsequently transduced with viral supernatants using spinfection. 48 hours after transduction, transduced cells were selected using puromycin for 5 days, and subsequently cultured in triplicates in 96 well plates. Cell viability was assessed at several time points after seeding using CellTiter-Blue (Promega). Cell viability was compared as percent viability of a corresponding condition using a shRNA directed against the luciferase gene.

### Orthotopic xenograft studies

We injected 200,000 BT16 cells into the forebrain (2 mm lateral and 1 mm anterior to bregma) of Crl:CD1-Foxn1^Nu/Nu^ nude mice (male and female, 6 to 8 weeks old). Mice were randomized to treatment arms. Both vehicle and abemaciclib treatment was scheduled daily at a concentration of 75 mg/kg by oral gavage for 3 weeks, with 2 day drug holidays on weekends. Animals were closely monitored.

### Western blotting

Whole cell lysates were prepared using RIPA buffer. Protein gels were run using 10% BOLT Bis-Tris or 4-12 % NuPAGE Bis-Tris polyacrylamide precast gels. Proteins were blotted onto PVDF membranes and detected using antibodies to CDK4, CDK6, cyclin D1, cyclin D2, cyclin D3, and GAPDH (see KEY RESOURCES TABLE) as per standard methods. Visualization was done using HRP-coupled, species-specific secondary antibodies.

### Immunohistochemical analyses

For immunohistochemistry to assess CDK6 and cyclin D2 in samples from ATRT patients, we used archived formalin-fixed, paraffin-embedded (FFPE) tumor material. FFPE tissue specimens were cut to 2,5 µm thick sections using a microtome, mounted on glass slides and subjected to immunhistochemical (IHC) staining. The stainings were performed using either CDK6 monoclonal rabbit anti-human antibody (Clone EPR4515, dilution 1:250, Abcam, Cambridge, UK) or Cyclin D2 monoclonal rabbit anti-human antibody (Clone D52F9, dilution 1:50, CellSignaling, Cambridge, UK). The IHC procedure was conducted using an automated immunostainer (BenchMark ULTRA IHC/ISH Staining Module, Hoffmann-La Roche, Basel, CH) together with the respective reagents (as listed below) and according to the protocols provided by the manufacturer. For detection the OptiView DAB IHC protocol was used including the following steps: Deparaffinization for 4 min at 72°C, washing with EZ Prep, incubation with Cell Conditioner No.1 for 64 min at 100°C, incubation with OV PEROX IHBTR for 4 min, incubation with primary antibody for 32 min at 37°C, and incubation in sequence with OV HQ UNIV LINKR, then incubation with OV HRP MULTIMER and then incubation with OV DAB and OV H2O2 for 8 min each, last incubation with OV Copper for 4 min. The counterstaining was performed with hematoxylin for 20 minutes upon which the tissues were incubated with BLUING REAGENT for 8 min prior to mounting the coverslips. Multiple intervening washing steps were included. All images were derived from tissue areas with evident tumor pathology, corresponding to approximately 1500 cells per sample and obtained using bright-field microscopy with a 40x objective (Olympus BX61). After adjusting a common threshold in all images to highlight stained cells, immunoreactivity was measured on ImageJ software as percentage area of CDK6 or cyclin D2 positive cells.

## QUANTIFICATION AND STATISTICAL ANALYSIS

No statistical methods were used to predetermine sample sizes except for *in vivo* studies, but the sample sizes here are similar to those reported in previous publications. Differences were considered statistically significant if *p* < 0.05, and sample sizes are indicated in each figure legend. For the comparison of gRNA read count distributions from CRISPR screens, we used the Kolmogorov-Smirnov test. For the analysis of CRISPR screens, two distinct metrics were used. One, we used the probability mass function of a hypergeometric distribution to assign *p*-values for each gene covered by the genome-wide gRNA library. Second, a negative binomial distribution (STARS algorithm) was used on a ranked list of gRNAs to assign the least probable perturbation for each gene as the STARS score. Permutation testing of the list of perturbations was used to calculate FDR statistics as previously described (Doench et al., 2016). For all correlation studies, a line derived from nonlinear regression was fitted onto the data, and correlation was computed using Pearson’s *r*. To compare the distribution of z scores relating to drug screen data, we used the Wilcoxon signed-rank test and calculated the corresponding effect size. Intersection analyses for genetic dependencies were calculated using the *SuperExactTest* package (Wang et al., 2015). All growth rate inhibition assays were analyzed using the *GRmetrics* package (Hafner et al., 2016). For survival analyses, we used the Log-rank test, and survival was defined as the time period between surgery and the onset of neurological symptoms, or the presence of any other exclusion criteria defined by local authorities. Differential gene expression analyses from RNA-seq data was performed using DESeq2 (Love et al., 2014), using a cutoff of log_2_ fold change > 1 and adjusted *p*-value < 0.01 for significance. For comparisons of more than 2 groups across more than 2 variables, we used Two-way ANOVA with Dunnett correction for multiple testing. All statistical analyses were performed in GraphPad Prism 8 or R.

---

**Figure S1.**
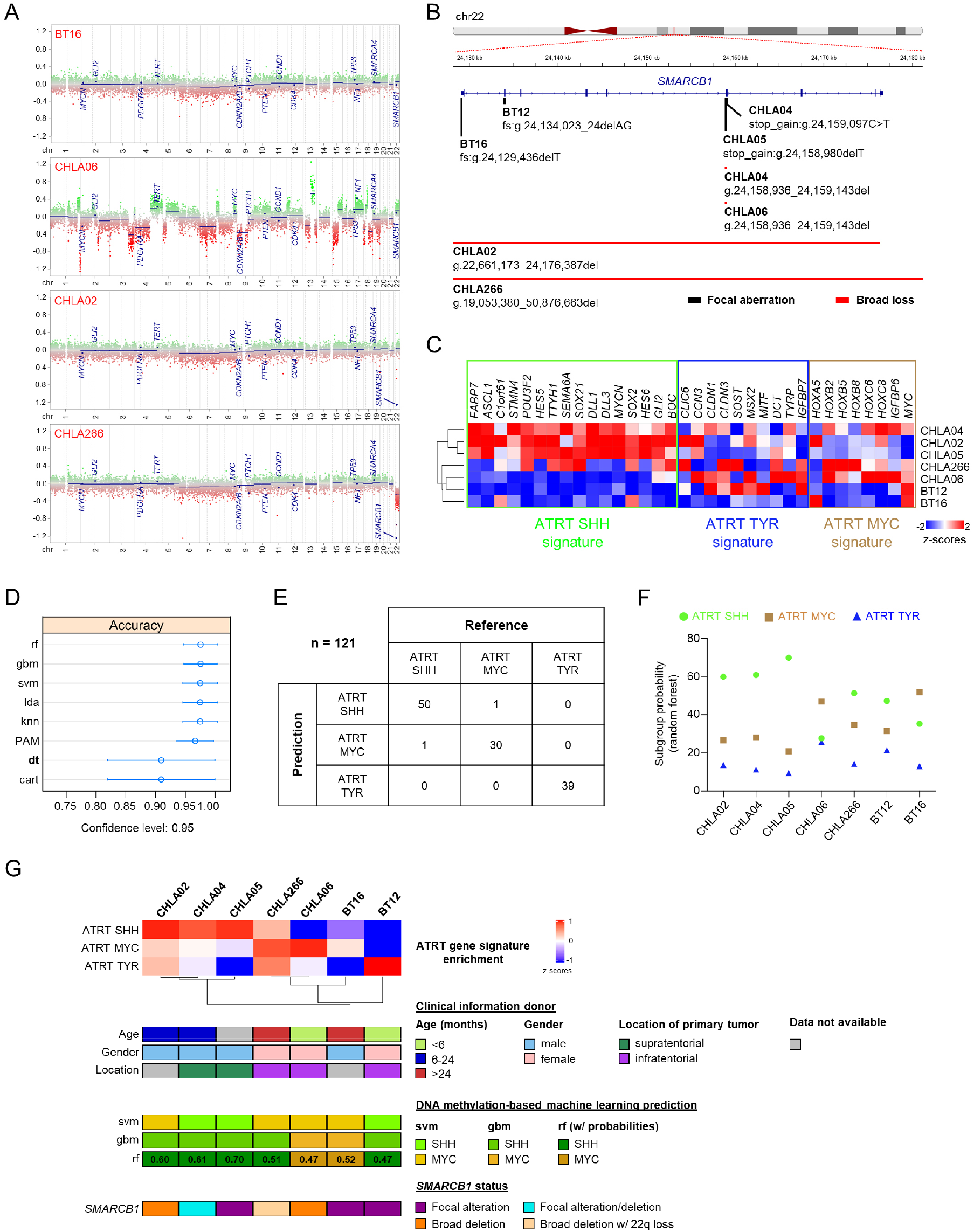
Detailed molecular classification of human ATRT cell lines, related to Figure 1. (A) Representative CNV plots for selected ATRT cell lines. Brain-tumor associated genes are highlighted. (B) Overview of *SMARCB1* alterations in ATRT cell lines. (C) Heatmap illustrating the enrichment of ATRT subgroup-associated gene signatures in ATRT cell lines. (D) Accuracy of several machine learning algorithms in predicting ATRT subgroups in the validation cohort. (E) Confusion matrix illustrating accuracy of the random forest model in predicting ATRT subgroups in the test cohort. (F) Subgroup probabilities for human ATRT cell lines as predicted by the random forest model. (G) Overview of key molecular and clinical parameters of ATRT cell lines including prediction of subgroup affiliation based on gene expression and global DNA methylation.

**Figure S2.**
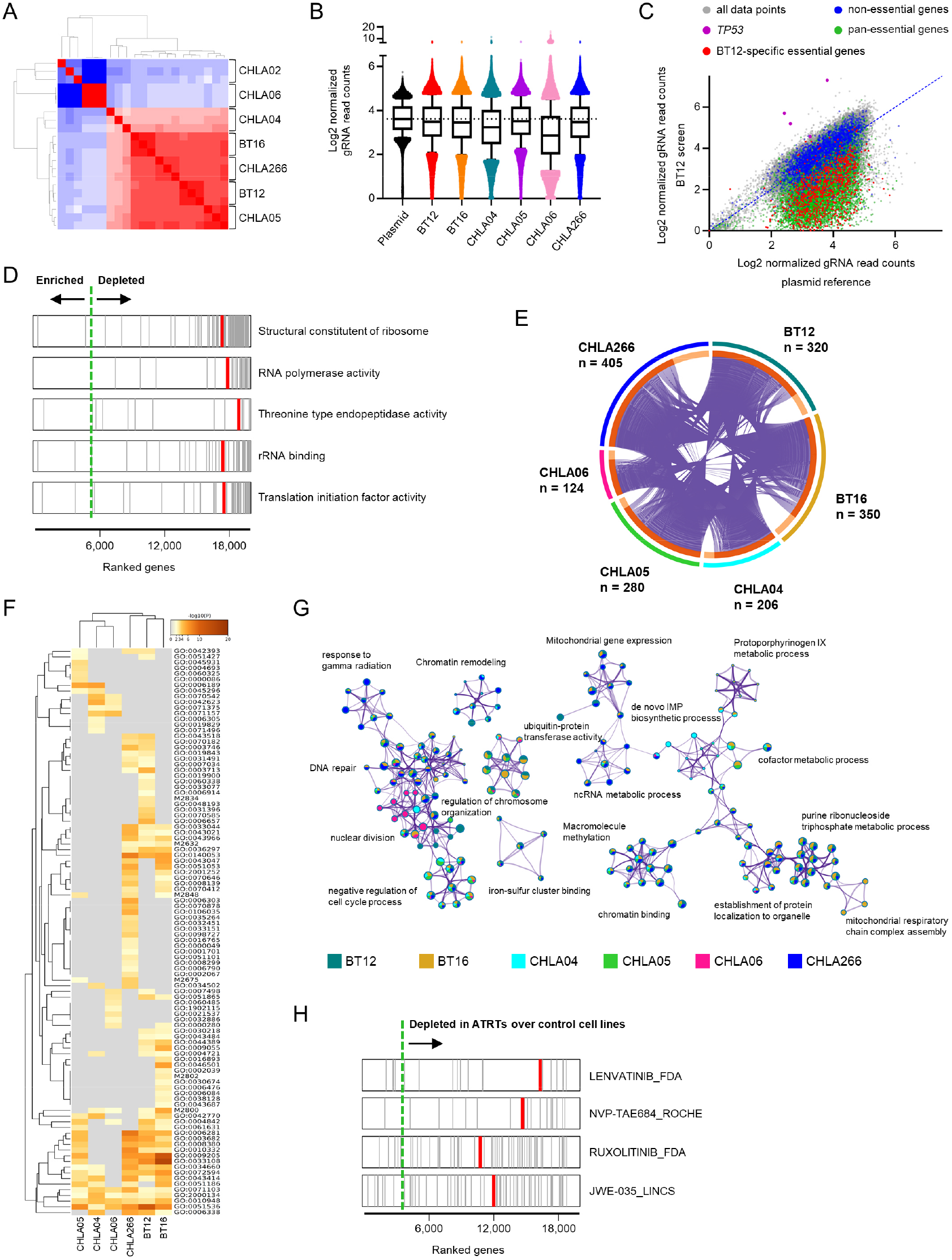
Detailed molecular classification of human ATRT cell lines, related to Figure 1. (A) Heatmap showing pairwise correlations (Pearson’s r) of pseudo-replicates from CRISPR/Cas9 ATRT screens based on normalized gRNA read counts. (B) Plotting of normalized gRNA read counts for plasmid reference and 6 ATRT CRISPR/Cas9 screens. (C) Pairwise correlations of normalized gRNA read counts from the reference control and the BT12 CRISPR/Cas9 screens, highlighting non-essential and pan-essential genes. Also, significantly enriched and cell context-specific depleted gRNAs are color-coded as well. (D) GSEA analysis showing the top 5 most significantly depleted gene sets averaged across 6 ATRT CRISPR/Cas9 screens. (E) Circos plot illustrating the overlap of cell context-specific dependencies from 6 distinct ATRT cell lines. (F) Heatmap of the top 100 most significantly enriched gene ontology terms in context-specific dependencies from ATRT cell lines. (G) Network layout of representative gene ontology terms affected in CRISPR/Cas9 screens, with circle size being proportional to number of input genes, and circle color indicating contributing ATRT cell lines. (H) Enrichment of selected drug signatures using the drug signatures database in ATRT cell line dependencies as compared to non-ATRT dependencies. To construct a ranked list of gene dependencies, depletion of enrichment of a specific gene was determined by contrasting results from 6 ATRT screens with results from essentiality screens using 4 non-ATRT cell lines (A375, HAP1, Meljuso, Ovcar8).

**Figure S3.**
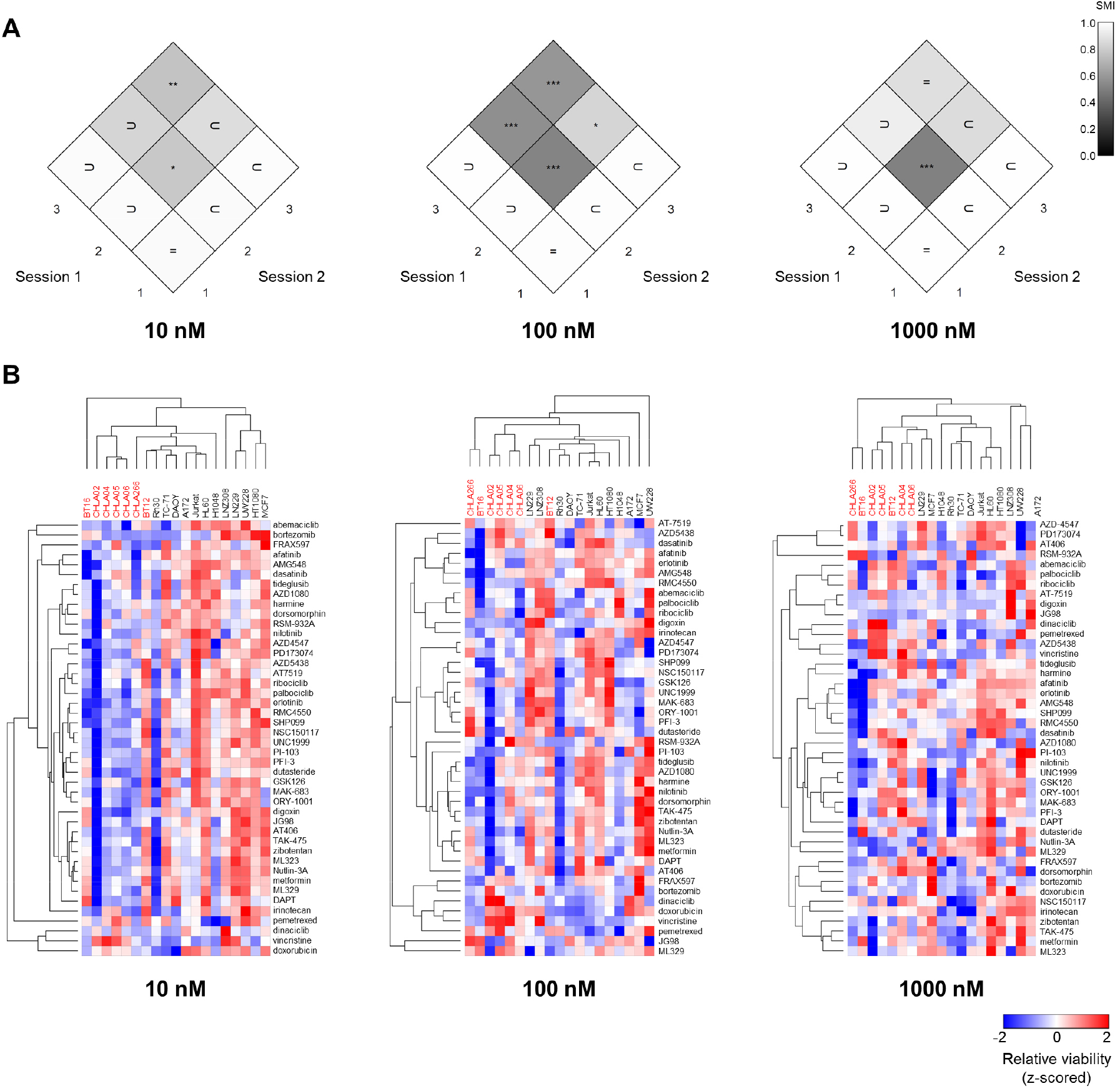
Extended analyses for 3 dose drug screen, related to Figure 3. (A) Diamond plots showing the similarity matrices index for the comparison of session 1 and session 2 of the 3 dose drug screen based on the effects on cell viability relative to DMSO control. (B) Heatmaps illustrating the z-scored effect of the ATRT drug library on cell viability as compared to DMSO control (1 minus Pearson correlation, average linkage). ATRT cell lines are indicated in red.

**Figure S4.**
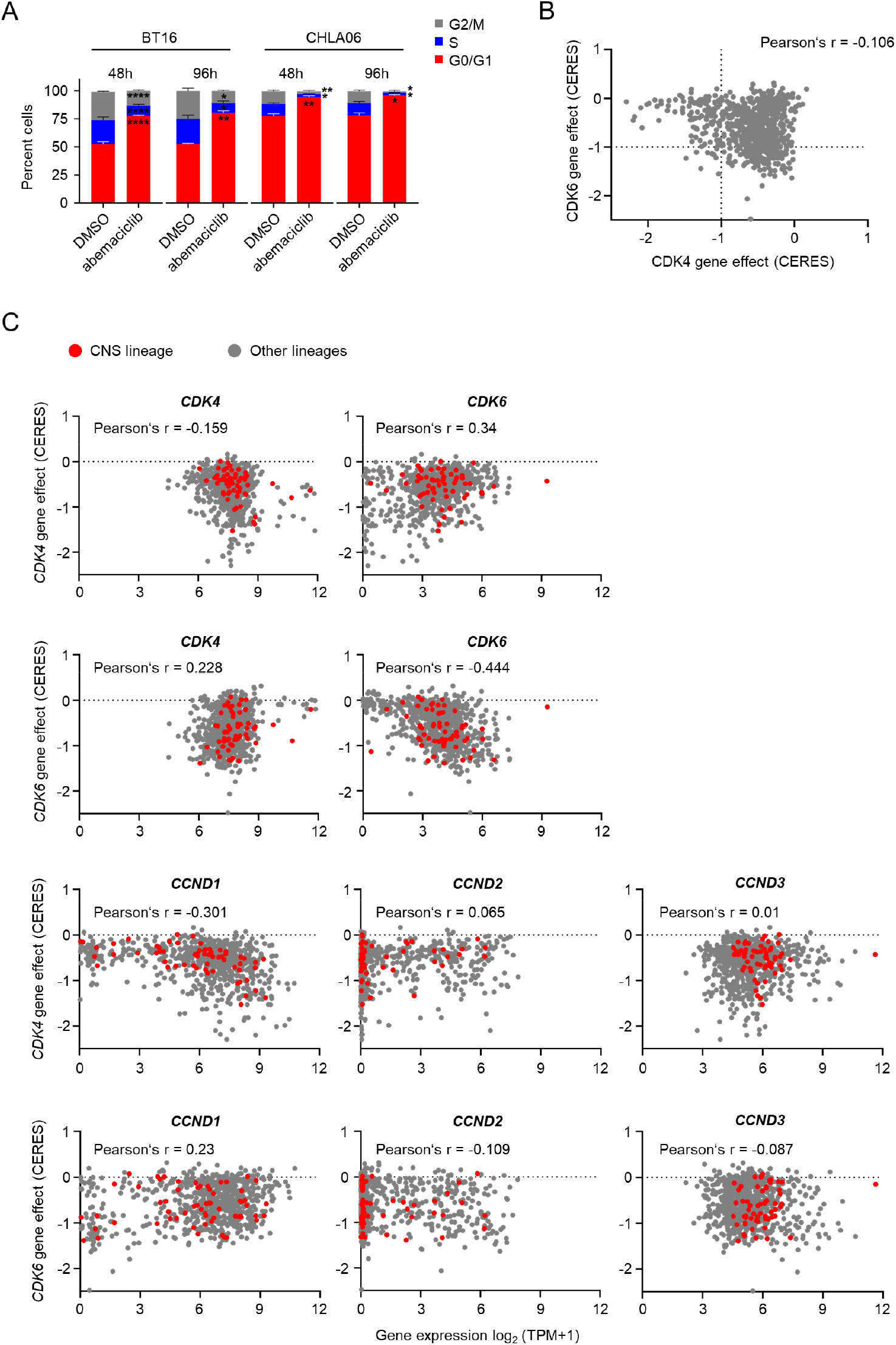
Extended analyses for CDK4/6 inhibitor sensitivity in ATRT cells, related to Figure 5. (A) FACS analyses using PI staining illustrating cell cycle phase shift after abemaciclib treatment in ATRT cell lines. (B) Correlation of *CDK4* and *CDK6* dependency (CERES scores) for a total of 789 human cancer cell lines from the Achilles project (https://depmap.org/portal). (C) Correlation of *CDK4* and *CDK6* dependency and mRNA expression of *CDK4, CDK6, CCND1, CCND2*, and *CCND3* in 789 human cancer cell lines. Central nervous system cancer cell lines are indicated in red.

**Figure S5.**
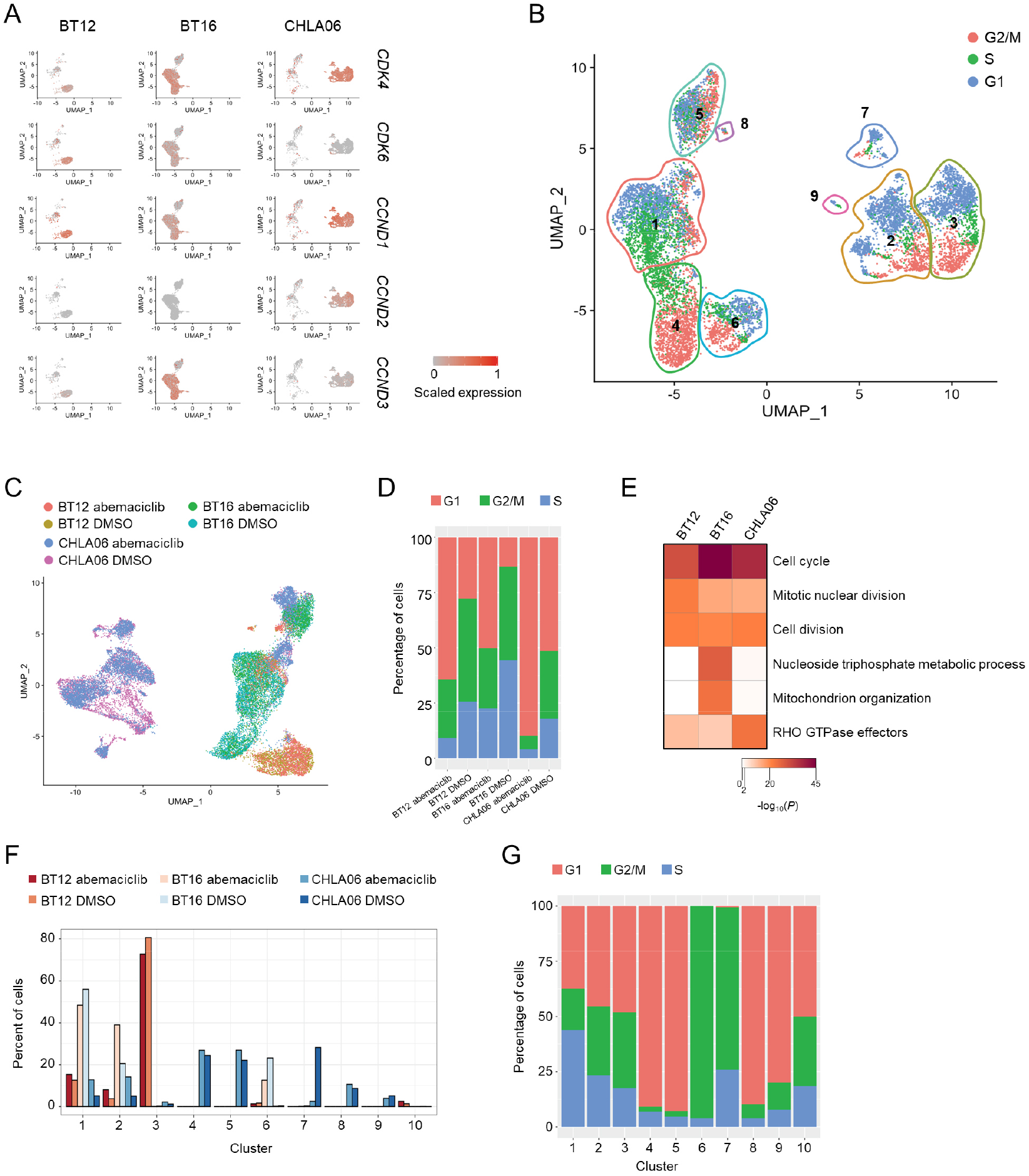
Single cell RNA-seq data of human ATRT cell lines in the context of *CDK4* and *CDK6* dependency, related to Figure 6. (A) UMAP embedding of vehicle-treated ATRT cell lines (BT12, BT16, and CHLA06) with projected expression of *CDK4, CDK6, CCND1, CCND2*, and *CCND3*. (B) UMAP embedding of scRNA-seq data from control BT12, BT16, and CHLA06 ATRT cells with cell cycle phase prediction based on cell cycle marker genes. Corresponding 9 transcriptionally distinct cell clusters are indicated according to Figure 6E. (C) UMAP plot of ATRT cell lines treated with vehicle or abemaciclib color-coded by cell line and treatment condition. (D) Bar graph showing the percentage of cells per cell line and condition in indicated phases of the cell cycle. (E) Heatmap showing the top 3 enriched process networks per cell line by abemaciclib treatment as compared to vehicle. (F) Percentage of cells from abemaciclib- and vehicle-treated BT12, BT16, and CHLA06 cells per cluster from corresponding cell line and treatment condition. (G) Bar graph illustrating percentage of cells from distinct clusters in indicated phases of the cell cycle.

